# Individual Differences in Dopaminergic Modulation of Exploration-Exploitation Behaviour

**DOI:** 10.64898/2026.01.30.702919

**Authors:** Elke Smith, Hendrik Theis, Thilo van Eimeren, Kilian Knauth, Deniz Tuzsus, Angela Brands, David Mathar, Jan Peters

**Affiliations:** Department of Psychology, Biological Psychology, University of Cologne, Cologne, Germany; Department of Nuclear Medicine, Faculty of Medicine and University Hospital Cologne, University of Cologne, Cologne, Germany; Department of Neurology, Faculty of Medicine and University Hospital Cologne, University of Cologne, Cologne, Germany; Department of Psychiatry and Psychotherapy, University of Bonn, Germany

**Keywords:** dopamine, L-DOPA, reinforcement learning, exploration-exploitation, interindividual differences

## Abstract

Dopamine (DA) has been implicated in exploration-exploitation behaviour, i.e., exploring novel, potentially better options vs. exploiting known, previously rewarding options. Impairments in this trade-off occur in psychiatric disorders involving DAergic dysfunction, including addiction and schizophrenia. Pharmacological studies revealed a contribution of DA to exploration, but inconsistent findings suggest that interindividual variability in baseline DA may modulate effects. To address this, we investigated the effects of the DA precursor L-DOPA on exploration-exploitation during reinforcement learning in a sample of N = 75 healthy participants (n = 32 women), following a randomised, double-blind, placebo-controlled, pre-registered design (https://osf.io/p2r7u). We assessed whether putative baseline DA markers, including spontaneous eye blink rate, working memory (WM) capacity, and impulsivity, modulated drug effects and probed visual fixation patterns and pupil dilation as markers of exploration. L-DOPA had no overall effect on computational model parameters of random exploration, directed exploration or choice perseveration. WM capacity moderated drug effects on random exploration, with stronger effects at higher WM capacity. Remaining DA proxies showed no credible effects. Pooling the data from male participants with that from an earlier male-only study (Chakroun et al., 2020; total N = 74), L-DOPA increased uncertainty-dependent value weighting and perseveration strength, while decreasing habit updating, indicating a stronger tendency to repeat previous choices and slower decay of their influence over time. No credible drug effects were observed in female participants. Pupil dilation was tonically increased under L-DOPA and scaled with exploration behaviour and prediction error, confirming that pupillometry can index exploration-exploitation dynamics. Visual exploration patterns reflected uncertainty-driven sampling, but were unaffected by L-DOPA. Taken together, results suggest that DAergic modulation of exploration and perseveration behaviour may be contingent on cognitive capacity and sex, rather than exerting uniform effects across individuals.

## 1 INTRODUCTION

Decision-making involves choosing between familiar options with known rewards and novel options with uncertain but potentially higher rewards, a dilemma known as the exploration-exploitation trade-off (Cohen et al., 2007; Sutton et al., 1998). A real-world example is the decision between staying in a known, secure job (exploitation) versus exploring new career paths (exploration). Finding the right balance is crucial for reward maximisation in dynamic environments. Exploration can improve long-term outcomes, but excessive exploration may reduce reward accumulation due to suboptimal switching. Conversely, excessive exploitation may lead to individuals sticking to suboptimal policies. Reinforcement learning (RL) is a framework for modelling decision-making through trial and error, balancing exploratory and exploitative actions to maximise cumulative reward over time (Sutton et al., 1998). Early theories, such as Thorndike’s Law of Effect, laid the foundation for RL by linking rewards to repeated actions (Thorndike, 1927), which is now widely applied to study human decision-making (Dayan & Daw, 2008; Lai & Gershman, 2024; Wise et al., 2024), and to develop machine learning systems in domains such as robotics and autonomous systems (Thompson, 2024).

RL algorithms adapt through feedback, where the neurotransmitter dopamine (DA) plays a key role by signalling reward prediction errors (RPEs), i.e., the discrepancy between anticipated and actual rewards to inform future choices (Glimcher, 2011; Schultz, 2016, 2024). DAergic activity in the striatum and prefrontal cortex modulates the strength of associations between actions and outcomes (Guitart-Masip et al., 2012; Hart et al., 2024; Jocham et al., 2011). Importantly, DA’s role extends beyond encoding RPEs and reinforcing associations – it also contributes to goal-directed behaviour (Balleine & O’Doherty, 2010; Gershman & Uchida, 2019; Goto & Grace, 2005) and flexible decision-making when balancing exploration and exploitation (Borwick et al., 2020; Chakroun et al., 2020).

Conceptually, exploration is commonly further divided into random and uncertainty-based exploration. Random exploration reflects choice randomisation (Daw et al., 2006; Wilson et al., 2014, 2021), often modelled by the slope of the softmax action selection function (Gershman, 2018; Sutton & Barto, 2018). In contrast, directed exploration entails the strategic selection of uncertain options for information gain (Cohen et al., 2007; Wilson et al., 2014, 2021). Random and directed exploration can therefore be conceptually linked to the common model-free vs. model-based distinction in action selection, where only model-based decisions (as in directed exploration) involve an explicit model of the environment (Balleine & O’Doherty, 2010; Daw et al., 2011). DA is thought to affect both systems, reinforcing actions based on past rewards in model-free learning (Daw & Tobler, 2014; Shohamy & Daw, 2014) and supporting flexible updating of internal models in model-based learning (Akam & Walton, 2021; Gershman & Uchida, 2019; Taira & Sharpe, 2025).

Understanding the role of DA in this trade-off is crucial for unraveling the fundamental mechanisms underlying human decision-making, and is also of clinical relevance. DA dysregulation, beyond Parkinson’s disease (Lotharius & Brundin, 2002), is linked to addiction (Trifilieff et al., 2017), depression (Dunlop & Nemeroff, 2007) and ADHD (MacDonald et al., 2024), all of which involve maladaptive changes in flexible decision-making (Ersche et al., 2008; Rupprechter et al., 2018; Shook et al., 2005). Although exploration-exploitation may be a candidate transdiagnostic mechanism underlying depressive, anxiety, and psychotic disorders, inconsistent findings warrant further investigation (Lloyd et al., 2024). Nevertheless, across disorders, imbalances in DAergic and noradrenergic pathways and disrupted representations of uncertainty appear to be key mechanisms driving maladaptive decision-making (Jami et al., 2025).

Goal-directed behaviour relies on uncertainty signals to guide choices towards lesser-known, i.e., more informative options, with the rostrolateral prefrontal cortex playing a key role by tracking relative uncertainty (Badre et al., 2012; Culbreth et al., 2023). DA levels have been linked to both increases and reductions in exploration. DA transporter blockade (DAT) in monkeys increased novelty-seeking behaviour (Costa et al., 2014). Similarly, in humans, the DA precursor L-DOPA increased random exploration in a two-step RL task (Kroemer et al., 2019). Also, the Catechol-O-methyltransferase (COMT) inhibitor tolcapone, which enhances DA availability primarily in the prefrontal cortex, increased uncertainty-driven exploration in humans (Kayser et al., 2015).

On the other hand, decreasing DA neurotransmission in rats using flupenthixol, a D1/D2 receptor antagonist, increased random exploration (Cinotti et al., 2019). Also, increased choice randomness was found when reducing DA transmission in mice (via genetic disruption of NMDA and glutamate receptors) and in hyperdopaminergic dopamine-transporter knockdown mice (Beeler et al., 2010; Cieślak et al., 2018). Similarly, higher serum levels of amisulpride, a D2/D3 receptor antagonist, were associated with increased random exploration in humans (Mikus et al., 2022). In a rodent study, increasing DA activity with apomorphine was found to decrease exploration, while decreasing DA activity with flupenthixol resulted in increased exploration, and modelling suggesting that DA modulated exploration behaviour in rats via an upregulation of decision noise (Chen et al., 2024). However, other studies have suggested more nuanced effects. For example, L-DOPA attenuated directed exploration in a four-armed restless bandit task, but did not reliably affect random exploration or exploitation (Chakroun et al., 2020). Also, patients with Parkinson’s disease off medication showed increased directed exploration but no changes in random exploration (Meder et al., 2025). Of note, in some Parkinson’s disease patients, dopamine agonists and L-DOPA have been linked to impulse control disorders, reflecting altered decision-making (Joutsa et al., 2012; Voon et al., 2017). The noradrenaline (NA) system has also long been implicated in exploration (Aston-Jones & Cohen, 2005). Studies show that lower NA levels are associated with less random exploration and more exploitation (Dubois et al., 2021; Turner et al., 2024). Another study reported a partly different pattern, where lower NA reduced the use of value information, but increased switching after high rewards (Cremer et al., 2023).

Inconsistencies in DA effects on the exploration-exploitation trade-off may stem from differences in experimental design, task types, and DA-modulating substances (e.g., flupenthixol, amisulpride, L-DOPA). These drugs differentially interact with different receptors (e.g., D1 vs. D2). Furthermore, exploration is differentially operationalised across studies. DA’s influence on exploration may also depend on the type of strategy, i.e., directed vs. random (Chakroun et al., 2020). Further, mixed findings may be explained by an inverted U-shaped relationship between DA and cognitive function (Cools & D’Esposito, 2011; Gjedde et al., 2010; Petzold et al., 2019). In this model, increasing DA levels may improve or impair performance depending on individual baseline levels. We did not observe such an association for temporal discounting (Smith et al., 2025), but the proposed relationship may still apply to exploration-exploitation behaviour.

To address these issues, we conducted a placebo-controlled, double-blind pharmacological intervention using the DA precursor L-DOPA to assess changes in exploration-exploitation behaviour and the role of individual differences in putative proxies of baseline DA, including spontaneous eye blink rate (sEBR) (Cools & D’Esposito, 2011; Jongkees & Colzato, 2016; Kaminer et al., 2011), working memory (WM) capacity (Braun et al., 2021; Cools & D’Esposito, 2011; Matzel & Sauce, 2023) and impulsivity (Kayser et al., 2012; London, 2020). Data from a temporal discounting task from this study were recently published (Smith et al., 2025). After a baseline screening for DA proxies, participants completed a four-armed restless bandit task (Chakroun et al., 2020; Daw et al., 2006; Wiehler et al., 2021) under L-DOPA or placebo (counterbalanced, within-subjects), while visual fixations and pupil dilation were measured as markers of explore-exploit tendencies (Jepma & Nieuwenhuis, 2011; Stojić et al., 2020; Van Slooten et al., 2018). We used hierarchical Bayesian modelling to assess DA effects on random and directed exploration, as well as choice perseveration (Miller et al., 2019; Rutledge et al., 2009). We then examined the degree to which DA proxy measures modulated drug effects, both in our de novo mixed sex sample (*N* = 75, *n* = 32 female), and in a pooled sample of only male participants (Chakroun et al., 2020) to explore potential sex-specific effects.

### Hypotheses

As preregistered (see https://osf.io/p2r7u), we predicted participants to show signatures of both exploitation and exploration during task performance (Chakroun et al., 2020; Daw et al., 2006; Speekenbrink & Konstantinidis, 2015; Wiehler et al., 2021), and predicted that a softmax model with a directed exploration term would best account for the data. Although we initially predicted L-DOPA to reduce response times (RTs) by increasing response vigour (Rihet et al., 2002; Westbrook et al., 2020), our recent analysis showed it increased RTs under L-DOPA in a temporal discounting task (Smith et al., 2025). In this mixed-sex sample, we predicted to replicate our previous finding of reduced uncertainty-based exploration under L-DOPA in men, with random exploration and perseveration unaffected (Chakroun et al., 2020). Based on the inverted U-shaped hypothesis of DA function (Cools & D’Esposito, 2011) suggesting that cognitive performance is optimal at intermediate DA levels and impaired at both low and high levels, a DAergic drug is expected to move individuals along the performance curve, Following this logic, we expected drug-induced changes in exploration–exploitation behaviour to vary linearly with baseline DA proxy measures. We also explored potential non-linear (quadratic) effects. We expected pupil dilation to decrease during exploitation and increase during exploration (Fan et al., 2023; Gilzenrat et al., 2010; Hayes & Petrov, 2016; Jepma & Nieuwenhuis, 2011). Further, L-DOPA was predicted to generally increase pupil dilation (Korczyn & Keren, 1982; Spiers & Calne, 1969). We also explored the degree to which drug-induced changes in exploration-exploitation behaviour were linked to changes in pupil dilation and fixation patterns.

## 2 METHODS

### 2.1 Sample

#### 2.1.1 Sample 1 (de novo dataset, *N* = 75, mixed sex)

All study procedures were approved by the local ethics board of the Faculty of Medicine of the University of Cologne, and all participants provided informed written consent prior to participation. Recruitment of participants was carried out at the University of Cologne through university bulletins, mailing lists, and word-of-mouth referrals. Participants were included if they met the following criteria: normal weight as indicated by a body mass index (BMI) between 19-25 +/-2 (since the BMI does not account for body composition, Provencher et al. 2018), right-handedness, normal or corrected-to-normal vision, German as first language or profound German language skills, and hormonal contraception for women. General contraindications included the use of common or prescription medications, participation in other studies involving medication, alcohol or drug intoxication or abuse, significant vision impairment or strabismus, acute infections, pregnancy, high emotional stress or physical strain during the study period, past or current psychiatric disorders, neurological conditions, metabolic disorders, internal diseases, chronic pain syndrome, and complications from anesthesia. Specific contraindications related to L-DOPA intake included hypersensitivity to L-DOPA or benserazide, use of non-selective monoamine oxidase inhibitors, metoclopramide, or antihypertensive medications (e.g., reserpine), disorders of the central DAergic system (such as Parkinson’s disease), increased intraocular pressure (e.g., glaucoma), and breastfeeding.

Eighty-five participants were invited to participate in the study, of which nine did not complete the study (*n* = 3 due to side effects of L-DOPA intake such as nausea, and *n* = 6 for other reasons such as illness and organisational reasons). The data of one participant were excluded from all analyses, as we assumed that the person did not complete the task conscientiously. The participant exhibited many premature responses, and since this was the case for both testing sessions (mean RTs were 0.34 s and 0.04 s for session 1 and 2, and in 20% and 93% of the trials, respectively, RTs were below < 200 ms), we assumed that this reflects an attempt to complete the task as quickly as possible rather than a technical error. The final sample thus comprised *N* = 75 participants (all right-handed, *n* = 32 female), aged 25 to 40 (*M* = 28.29, *SD* = 3.20), having undergone *M* = 16.41 years of education (*SD* = 2.31, range = 10 to 18), with a BMI of *M* = 22.56 (*SD* = 1.91, range = 18 to 26). Scores for the BIS-15 (Meule et al., 2011; Spinella, 2007) ranged from 17 to 41 (*M* = 30.85, *SD* = 5.07). In the analysis of pupil dilation and the effects of the putative DA proxy measures on the drug effects (Bayesian regression model), *n* = 74 participants were included, since data for the BIS-15 and eye tracking data were missing for one participant due to technical issues. The sample has been described in Smith et al. (2025).

#### 2.1.2 Sample 2 (pooled sample, *N* = 74 men)

To reassess the drug effects on model parameters in a larger sample, we pooled the data of the male participants from sample 1 with those from Chakroun et al. (2020), who tested a European male-only sample on the same RL task. This resulted in a pooled male sample of *N* = 74 (our male participants plus *n* = 31 from Chakroun et al., aged 19 to 40). Female participants from sample 1 (*n* = 32) were analysed separately. For a detailed description of the original male sample, see Chakroun et al. (2020).

### 2.2 Study design

#### 2.2.1 Procedure

The study was conducted using a repeated-measures, within-subject design (see preregistration at https://osf.io/a4k9j/). After completing a medical assessment by a physician to screen for any contraindications (see section 2.1.1), eligible participants were invited to attend three separate testing sessions. In the first session, participants underwent a baseline screening, which included the recording of sEBR (using a webcam), assessment of WM capacity (digit span; Wechsler, 2008; listening span; van den Noort et al., 2008; operation span; Foster et al., 2015), psychometric assessment of impulsivity (Barratt Impulsiveness Scale, BIS-15; Meule et al., 2011; Spinella, 2007) and the collection of demographic information. After the baseline screening, the participants attended two identical double-blind placebo-controlled testing sessions during which they received either L-DOPA or placebo. These sessions were carried out at one-week intervals at a similar time of day, and for women not during the pill break. In some cases, the interval was either shortened or extended for health or organisational reasons. Participants were instructed to refrain from eating for at least 2 hours prior to testing. Before each drug session, they underwent a screening for pregnancy and the use of alcohol and drugs (including THC, cocaine, MDMA, amphetamines, benzodiazepines, barbiturates, methamphetamine, morphine/heroin, methadone, and tricyclic antidepressants) and excluded if tested positive.

Thirty minutes before testing (to ensure peak plasma concentration during the testing phase), participants were administered a tablet containing either 150 mg of L-DOPA (plus 37.5 mg benserazide) or a placebo (maize starch). Physiological parameters and well-being were assessed at the start, during the course, and at the end of each session to ensure participant safety throughout the experimental process. Following the waiting period, participants completed an intertemporal choice task (Smith et al., 2025), a RL task, and a visual pattern perception task (Smith et al., 2024). They were financially compensated for their participation, with an additional variable bonus based on the collected points (see section 2.2.2). For a description of the procedure undergone by participants in sample 2 (pooled sample), which involved the same 150 mg dose, we refer the reader to Chakroun et al. (2020).

#### 2.2.2 Reinforcement learning task

We adapted the four-armed restless bandit task from Daw and colleagues (2006). In light of modelling analyses showing adequate parameter recovery for 200 trials (Danwitz et al., 2022), we reduced trial numbers to 200 compared to our previous study (Chakroun et al., 2020) and the original paper (Daw et al., 2006) that used 300 trials. Participants were instructed that 5% of the points earned (5 cents per 100 points) in the experiment would be paid out at the end. Upon completion of the task, the sum of points earned was displayed on the screen. Before testing, the participants were familiarised with the task by performing a training phase of 10 trials. The practice trials were followed by a 5-point binocular calibration for accurate gaze tracking. The task was divided into four blocks of 50 trials, interleaved by short breaks. A trial started with the display of four squares of different colours (“bandits”) to choose between. Since pupil dilation is affected by luminance, we controlled the luminance of the stimuli. All bandits had the same mean intensity value and were presented against an isoluminant grey background. The mean payoff linked to a bandit varied randomly across trials following a decaying Gaussian random walk. Two different, pseudorandomly assigned random walks were used for the placebo and L-DOPA sessions. A bandit was selected by button press (“I”, “O”, “K” or “L” of the keyboard) using the right hand, with an RT limit of 1.5 s. If no button was pressed within this time window, a red “X” was displayed for 2.2 s at the centre of the screen to indicate a missing response with no earnings. Following choices within the response window, the selected option was highlighted (keeping overall luminance constant) for 2 s. Hereafter, the number of points earned in the trial was displayed for 1 s in white colour in the bandit’s centre. After the feedback period, all bandits were masked out, and a white fixation cross remained at the centre of the screen until the trial ended after 5 s. The trials were interleaved with inter-trial intervals (ITI) with random durations drawn from a linearly spaced vector between 2 and 3 s.

### 2.3 Analysis

#### 2.3.1 Model-agnostic trial classification and analysis (sample 1, *N* = 75)

Model-agnostic trial classification involved four key definitions. Stay and switch denote repeating and changing the previous choice, respectively. Uncertainty-based exploration was operationalised as choosing the bandit that had not been selected for the greatest number of consecutive trials. Random exploration was defined as switch trials that did not meet the criteria for uncertainty-based exploration. Since exploitation and perseveration cannot be disentangled by means of model-agnostic measures, we further focus on stay and switch proportions (i.e., repeating or changing the previous choice, respectively). Optimal choices referred to selecting the option with the highest value.

To assess differences in model-agnostic measures (total points, stay and switch decisions, random exploration, and optimal choices) between drug conditions, we report Bayes’ factors (Lee & Wagenmakers, 2014) calculated with JASP (JASP Team, 2024; version 0.19.0). Ensuing from the hypothesis of reduced uncertainty-based exploration and no change in random exploration, by implication, we expected an increase in the proportion of stay trials (one-sided test, choosing the same bandit as in the previous trial), and correspondingly, a decrease in the proportion of switch trials (one-sided test, choosing a different bandit), i.e., less stay trials under placebo compared to L-DOPA, and more switch trials under placebo compared to L-DOPA. Therefore, one-sided tests were used for response times, uncertainty-based exploration, stay and switch trials, while two-sided tests were used for total points, optimal choices and random exploration. We use Bayes’ factor (BF) notation to indicate evidence for the null versus alternative hypothesis: 0+/0– indicate evidence for the null in directional tests with positive/negative expected effects, +0/–0 indicate evidence for the alternative in directional tests with positive/negative expected effects, and 01/10 indicate evidence for the null or alternative, respectively, in two-sided tests.

#### 2.3.2 Reinforcement learning models

The computational models for the four-armed restless bandit task consist of two components: a learning rule, which updates estimates of an option’s subjective value, and a choice rule, which governs decisions based on those values. We compared two learning rules: the delta rule from temporal difference learning (Sutton & Barto, 2018) and a Bayesian learner (Daw et al., 2006) that accounted better for behavioural data in several previous studies (Chakroun et al., 2020; Wiehler et al., 2021a). Each will be paired with four choice rules based on the softmax selection rule, resulting in eight models. An overview of the models and their parameters is provided in Tables 1 to 3.

**Table 1.** Free and fixed parameters of the delta rule models.

|  | Parameter | Function |
| --- | --- | --- |
| Free | $\alpha$ | Learning rate |
| | $\beta$ | Random exploration (softmax inverse temperature) |
| | $\varphi$ | Directed exploration |
| | $\rho$ | Perseveration |
| Fixed | $v_1$ | Initial expected subjective value |

Note. The free parameters are only listed for the subject-level.

##### Delta rule

Following the delta rule, participants update the expected subjective value *v* of the chosen bandit based on the prediction error *δ*, which is the difference between the obtained reward *r* and the expected reward for trial *t* :

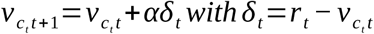

The parameter *α* represents the learning rate, a free parameter between zero and one, that controls the proportion of the prediction error used to update the Q-value. Q-values of unchosen bandits remain unchanged. Table 1 provides an overview of the model parameters.

##### Bayesian learner

The Bayesian learner model uses the Kalman filter algorithm (Kalman, 1960; Moore & Anderson, 1979) for updating reward values. It assumes that participants form an internal representation of the task’s true reward structure. At the start of each trial, participants have a prior belief about each bandit’s mean payoff, which is updated using Bayes’ theorem. The prior belief for bandit *i* on trial *t* is normally distributed with mean 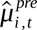 and 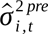. Following a choice, the chosen bandit’s prior is updated with the reward observation *r_t_*, resulting in a posterior distribution with updated mean 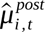 and variance 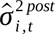 as follows:

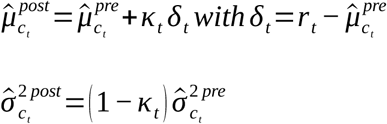

with

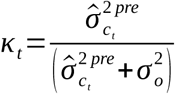

*κ* is the Kalman gain for trial *t*, which controls the proportion of the prediction error used to update the prior distribution. The Kalman gain varies from trial to trial, depending on the variance of the expected reward’s prior distribution 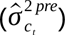 and the estimated observation variance 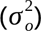 . The observation variance reflects the extent to which rewards fluctuate around the (estimated) mean reward of a bandit and indicates the reliability of a trial’s observed reward in estimating the true underlying mean reward. If the prior variance is large relative to the estimated variance (i.e., when a participant’s reward estimates are highly uncertain, but the reward observations are very reliable), the Kalman gain approaches one, and a large fraction of the prediction error is used to update the reward expectation. In contrast, when the prior variance is small relative to the estimated variance (i.e., when a participant’s reward estimates are very reliable, but the reward observations are highly noisy), the Kalman gain approaches zero, and only a small portion of the prediction error is used to update the reward expectation. The expected rewards (prior mean and variance) of the unchosen bandits are not updated within a trial, implying that their posterior distributions equal their prior distributions. Between trials, however, the prior distributions of all bandits are updated based on a participant’s assumption about the underlying Gaussian random walk as follows:

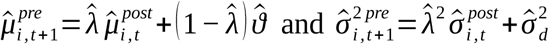

The updating process was initialised with the same prior distribution *N* 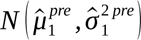, where 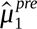 and 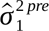 are treated as additional free model parameters.

##### Softmax choice rule

We implemented six variants of the softmax choice rule (Chakroun et al., 2020; McFadden, 1973; Sutton & Barto, 2018). The first and most basic variant only included the inverse temperature parameter *β*, reflecting choice stochasticity (random exploration). The probability *P_i,t_* of choosing bandit *i* on trial *t* was modelled as follows:

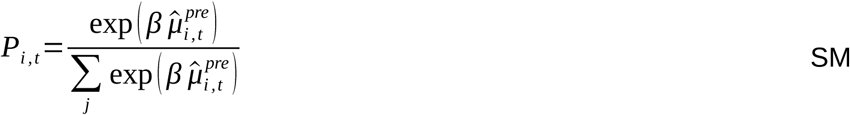

The second variant incorporated directed exploration by including a directed exploration bonus parameter *φ*, as implemented previously (Chakroun et al., 2020; Daw et al., 2006; Wiehler et al., 2021). The parameter *φ* reflects the degree to which choice probabilities are modulated by the uncertainty associated with a bandit:

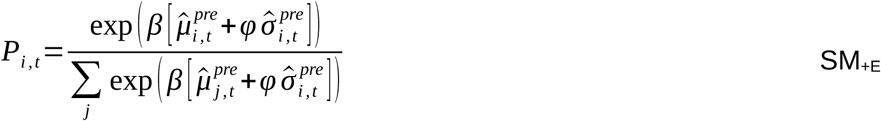

Depending on the learning rule, different approaches were used to quantify uncertainty. For the Bayesian learner, the uncertainty of an option was represented by the prior standard deviation of that option’s expected reward distribution 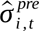. For the delta rule, we quantified uncertainty using two different counter-based metrics (Brands et al., 2025): (1) how many trials ago an option was chosen (uncertainty increases linearly with the number of trials, i.e. time passed, since an option was last chosen, referred to as “recency” heuristic in the following), and (2) how many alternative options have been chosen in the meantime (referred to as “diversity” heuristic in the following). The third choice rule variant included a first-order perseveration (FOP) parameter *ρ*, reflecting a bonus in form of a constant value added to the expected value of the previously chosen bandit (whereby *I* is an indicator function which equals 1 for the bandit chosen in the previous trial (indexed by *c_t−_*_1_), and 0 for the remaining unchosen bandits. This model corresponds to the best-fitting model as reported in Chakroun et al. (2020) and Wiehler et al. (2021).

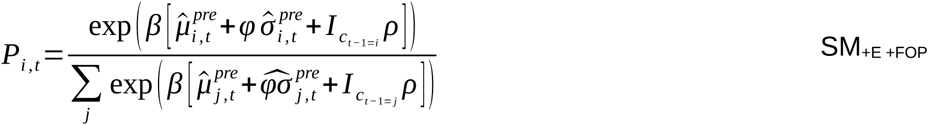

Extending our modelling approach beyond the preregistration, we additionally included a higher-order perseveration (HOP) term (Gershman, 2020; Lau & Glimcher, 2005; Miller et al., 2019), which we recently found to substantially improve model fit (Brands et al., 2025). Here, a habit strength vector *h* stores the choice history for each option and is updated trial-wise as follows:

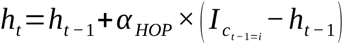

Again, *I* reflects an indicator function as described above. The degree to which habit strength *h_t_* is updated is controlled by the habit update parameter *α _HOP_* . *H* controls how quickly the influence of past choices updates (“habit update”, whereby higher values of *α _HOP_* implies faster updating, i.e. recent choices dominate, while lower values of *α _HOP_* implies slower habit decay, i.e., older choices still influence choices, reflecting a longer temporal integration. This model thus includes FOP as special case when *α _HOP_*=1. This is incorporated in the computation of choice probabilities as follows:

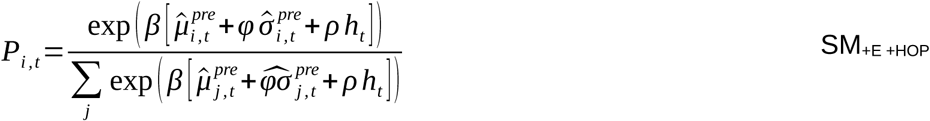

Here, *ρ* controls the degree to which habit strength *h_t_* influences current decisions (“perseveration strength”, whereby higher *ρ* implies a stronger incorporation of habit strength). The last choice rule variant included an additional total (summed across all bandits) estimated uncertainty (*Σσ*) term (Gershman, 2018; Gershman & Tzovaras, 2018). This term specifically modulates how strongly value estimates influence choice probabilities based on the agent’s overall uncertainty across all options:

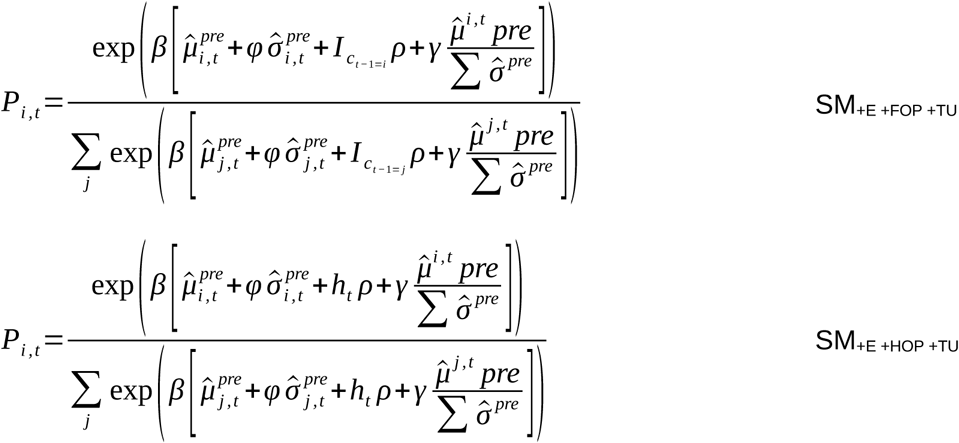

When total uncertainty is high (large *Σσ*), the denominator increases, reducing the magnitude of the value term (*μ* / *Σσ*). This diminishes the influence of value estimates on choice, allowing other decision components (exploration, perseveration) to drive choices more strongly. When total uncertainty is low (small *Σσ*), the denominator decreases, amplifying the value term (*μ* / *Σσ*). This increases the influence of value estimates on choice, promoting more value-driven (exploitative) decisions when uncertainty about option values is low. A positive *γ* would indicate that participants weight value estimates more strongly when uncertainty is low and less when uncertain is high. A negative *γ* would indicate the opposite pattern, i.e., stronger reliance on value estimates when uncertainty is high. For a summary of model parameters, see Table 2.

**Table 2.** Free and fixed parameters of the Bayesian learner models.

|  | Parameter | Function |
| --- | --- | --- |
| Free | $\beta$ | Random exploration (softmax inverse temperature) |
| | $\varphi$ | Directed exploration |
| | $\rho$ | Perseveration strength |
| | $\alpha_{HOP}$ | Habit update |
| | $\gamma$ | Total uncertainty weight |
| Fixed | $v_1$ | Initial expected subjective value |
| | $\hat{\lambda}$ | Decay |
| | $\hat{\vartheta}$ | Decay centre |
| | $\hat{\sigma}_o^2$ | Observed variance |
| | $\hat{\sigma}_d^2$ | Diffusion variance |
| | $\hat{\mu}_1^{pre}$ | Initial mean of prior expected reward |
| | $\hat{\sigma}_1^{pre}$ | Initial standard deviation of prior expected reward |

Note. The free parameters are only listed for the subject-level. HOP: higher-order perseveration.

**Table 3.** Model comparison for the condition-specific delta rule and Bayesian learner models. Comparison by LOO information criterion (estimate and standard error) and ELPD difference (estimate and standard error) with respect to the top model, ranked by predictive accuracy from highest to lowest.

| Model |  | Condition |  |  |  |  |  |  |  |
| --- | --- | --- | --- | --- | --- | --- | --- | --- | --- |
|  |  | Placebo |  |  |  | L-DOPA |  |  |  |
| Learning rule | Choice rule | LOO IC | SE | ELPD $\Delta$ | SE $\Delta$ | LOO IC | SE | ELPD $\Delta$ | SE $\Delta$ |
| Delta rule | SM<sub>+E</sub> +FOP (recency) | 23,082 | 675 | 0.0 | 0.0 | 22,524 | 746 | 0.0 | 0.0 |
|  | SM<sub>+E</sub> +FOP (diversity) | 23,343 | 663 | -130.0 | 43.0 | 22,754 | 746 | -115.2 | 31.2 |
|  | SM<sub>+E</sub> (recency) | 23,974 | 636 | -446.1 | 68.7 | 23,070 | 724 | -273.2 | 49.9 |
|  | SM<sub>+E</sub> (diversity) | 24,209 | 677 | -563.1 | 82.2 | 23,293 | 735 | -384.6 | 56.6 |
|  | SM | 24,545 | 683 | -731.3 | 102.3 | 23,651 | 754 | -563.9 | 98.2 |
| Bayesian learner | SM<sub>+E</sub> +HOP +TU | 22,452 | 674 | 0.0 | 0.0 | 21,716 | 752 | 0.0 | 0.0 |
|  | SM<sub>+E</sub> +HOP | 22,727 | 702 | -137.3 | 55.5 | 21,985 | 754 | -134.4 | 31.2 |
|  | SM<sub>+E</sub> +FOP +TU | 22,800 | 670 | -173.6 | 28.5 | 22,164 | 744 | -223.9 | 35.0 |
|  | SM<sub>+E</sub> +FOP | 23,073 | 709 | -310.1 | 72.3 | 22,426 | 748 | -354.9 | 51.6 |
|  | SM<sub>+E</sub> | 23,757 | 678 | -652.5 | 84.5 | 22,829 | 737 | -556.5 | 61.6 |
|  | SM | 24,430 | 699 | -988.7 | 112.0 | 23,644 | 781 | -964.2 | 144.7 |

Note. Model comparison was based exclusively on data from sample 1 (de novo dataset, $N = 75$ , mixed sex). LOO: leave-one-out; IC: information criterion; SE: standard error; ELPD: expected log pointwise predictive density; $\Delta$ : difference, SM: softmax; E: exploration; FOP: first-order perseveration; HOP: higher-order perseveration; TU: total uncertainty. Lower LOO IC scores indicate a better model fit. For the delta rule models, uncertainty was quantified using the counter-based metrics “recency”, i.e., time passed since an option was last chosen, and “diversity”, i.e., how many alternative options have been chosen in the meantime, respectively.

#### 2.3.3 Model comparison and drug-effect model

We first implemented separate models for each drug session to test the degree to which model comparison was affected by the drug. Following parameter estimation (see section 2.3.5), we compared the models’ predictive accuracy using leave-one-out (LOO) cross-validation based on Pareto-smoothed importance sampling (PSIS; Vehtari et al., 2017). The model with the highest predictive accuracy identified by model comparison (Bayesian learner with SM +E +HOP +TU) was then implemented as a full model including drug-induced changes in each parameter. In this model, the placebo condition was modelled as the baseline, and drug effects were modelled as additive condition-specific effects for each parameter.

#### 2.3.4 Regression model for DA proxy measures

To assess the relationship between DA effects and the putative DA proxy measures (BIS-15 scores, sEBR and WM capacity), we tested for a linear and non-linear (quadratic) relationship by means of Bayesian regression, regressing the drug effect parameters on the (squared) DA proxies. We used the DA proxy measures as individual regressors instead rather than a compound score, since the scores were only weakly correlated (−0.12 < *r* < 0.16). For WM capacity, we used the first principal component across all three WM tasks as a predictor (see supplementary Text S1, Table S2 and Figure S3). To provide an example, the regression model below outlines the relationship between the (squared) DA proxies and the drug effect on *φ* (directed exploration):

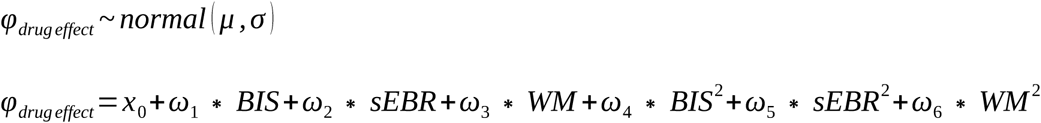

Here, *x*_0_ denotes the intercept, *ω*_1*−*3_ denote the regression coefficients for the DA proxy measures BIS-15 score (impulsivity), sEBR and WM capacity, respectively, and *ω*_4*−* 6_ denote the regression coefficients for the squared measures.

#### 2.3.5 Parameter estimation

Parameter estimation was conducted using Markov chain Monte Carlo (MCMC) as implemented in Stan (version 2.26.1) (Carpenter et al., 2017), R (version 4.3.1) (R Core Team, 2023) and the RStan interface (version 2.26.23) (Stan Development Team, 2023). All models were implemented as hierarchical models, in which single-subject parameters were drawn from group-level Gaussian distributions. We preregistered the use of Chakroun et al. (2020) posteriors as priors, but due to differences in modelling approaches (e.g., the inclusion of HOP and total uncertainty terms) we instead applied uniform priors over plausible ranges for all parameters (see supplementary Table S4). For the condition-wise models, sampling was performed with 10,000 iterations across 2 chains (8,000 warmup samples), for the drug-effect model, sampling was performed with 12,000 iterations across 2 chains (10,000 warmup samples). For the Bayesian regression models, sampling was performed with 2,000 iterations across 2 chains (1,000 warmup samples). Chain convergence for the single-subject and group-level parameters was assessed by inspecting the traces such that *R̂* ≤ 1.01 (Gelman & Rubin, 1992).

#### 2.3.6 Posterior predictive checks

To assess if the best-fitting model captures key patterns of the observed data, we conducted posterior predictive checks by comparing simulated to observed data across several behavioural metrics. For each trial and participant, we simulated 4,000 choices from each participant’s posterior distribution and compared them (averaged across simulations) to the observed choices. Since random walks were pseudorandomly assigned to the placebo and L-DOPA sessions, simulated and observed choices were compared separately for each random walk condition. We assessed predictive accuracy, defined as the proportion of trials where the model predictions matched the observed choices; the proportion of switches, defined as trials where participants chose a different option than on the previous trial; and the proportion of optimal choices, defined as selections of the currently highest-value option.

#### 2.3.7 Analysis of posterior distributions

We assessed the evidence for drug effects on model parameters by considering the highest posterior density intervals (HDIs) of the posterior distributions of the drug-effect parameters. Effects were considered reliable if zero was not included within the 95% HDI. For the regression model (see section 2.3.4) assessing a possible quadratic relationship between DA effects and the putative DA proxy measures, we also considered 95% HDIs.

#### 2.3.8 Visual fixation and pupil dilation

Preprocessing of visual fixation and pupil dilation data was conducted using MATLAB (version R2022a; The MathWorks, Inc., 2022). For both pupil dilation and gaze data, we excluded fast trials with a duration below 200 ms (Whelan, 2008). This decision was made because, given the sampling frequency of the eye tracker, trials with such short RTs would result in either no or very few samples during the pre-choice phase, making it invalid to draw valid inferences from them. These trials were also removed from the behavioural data for this analysis. As outlined in our preregistration, we intended to exclude data from participants with more than 90% of trials excluded (i.e., fast trials, no-response trials, and trials with many missing data points, as detailed below). However, this criterion was not met by any participant, and therefore, no participants were excluded.

##### Visual fixation

For fixation detection, we defined areas of interest (AOIs) based on the options’ borders, plus a tolerance of 1.5° visual angle. Fixations were defined as continuous binocular fixations with a minimum duration of 90 ms (Galley et al., 2015) falling within the respective AOI. We analysed fixations during the pre-choice phase only, i.e., from onset of the choice options until choice. Specifically, we assessed which bandit was fixated last before a choice was made and assessed the relationship between fixations and choices using a chi-square test. No-response trials and trials in which no option was fixated were not considered. Further, we calculated the number of fixation shifts, defined as changes of fixation from one option to another, and tested whether the number of fixation shifts was related to total uncertainty and uncertainty of the chosen option using a generalised linear mixed-effects model (GLMM) using MATLAB’s `fitglme` (using a Poisson distribution with a log link function). The model included drug condition, total uncertainty, and their interaction, as well as uncertainty of the chosen option and its interaction with drug, with participant included as a random intercept:

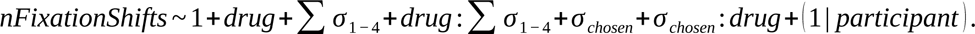

To assess the relationship between fixations and subsequent choices, we identified the last fixation prior to the decision and the corresponding choice made from the four options (excluding non-fixation and no-choice trials). A chi-squared test of independence was used to assess whether last fixations were predictive of the choices.

##### Pupil dilation

Prior to the analysis, clearly invalid (i.e., negative and zero values) samples were removed. Further, outliers, defined as data points exceeding the mean by more than 2 standard deviations, were removed from the time series. Trials with over 12% missing data points were excluded. Remaining missing data points (e.g., resulting from blinks) were linearly interpolated. The data were segmented as follows: (1) pre-stimulus baseline, i.e., 1 second time-window before the onset of the choice options until choice onset, (2) pre-choice phase, i.e., from onset of the choice options until choice, and (3) feedback, i.e., at onset of the reward feedback until the end of the feedback phase. For standardisation, we applied baseline correction using divisive baseline correction (corrected diameter = diameter during stimulus presentation / mean diameter during baseline window). We defined the last second of the preceding ITI as the pre-stimulus baseline. Since the pre-stimulus baseline before the first trial was insufficient, we used the mean of all subsequent baseline windows for correction. As a last step, the pupil data were downsampled to a frequency of 20 Hz. For final analysis, we used the mean pupil diameter across both eyes. To analyse effects of drug, time, decision variables and choice type on pupil diameter, we specified a GLMM using MATLAB’s `fitglme` (using an identity link function) as follows:

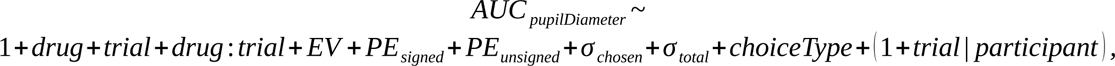

with the area under the curve (AUC) of the pupil diameter trajectory for each trial phase (pre-choice, feedback, and inter-trial interval) as dependent variable. AUC values were calculated using the trapezoidal numerical integration (`trapz` function in MATLAB) of the mean pupil diameter over time and normalised by the trial phase duration. The model included fixed effects for drug, since L-DOPA may cause general pupil dilation, trial number, and the interaction between drug and trial number, to test for potential drug-related changes in pupil responses over time (i.e., a decay across the course of the experiment). We also included random intercepts and slopes for each participant, allowing for individual variation in both baseline and the effect of trial progression. All continuous predictors (i.e., uncertainty measures and decision variables) were standardised. We ran separate models for all task phases (pre-choice, feedback and inter-trial interval). Decision variables were included as fixed effects, specifically, prediction error (signed and unsigned), expected value, uncertainty of chosen option (*σ _chosen_*), total uncertainty (*Σσ*), and choice type (random and directed exploration vs. exploitation coded as baseline), to examine how they related to pupil dilation. We classified trials as explore/exploit trials based on the model parameters for directed and random exploration, and exploitation, respectively. The model computes choice probabilities for each of the four options based on these parameters. The trial classification scheme aligns with the model parameters, such that choices were classified as exploitation when the option with the highest expected value was selected, as directed exploration when the option with the highest directed exploration bonus was chosen, and as random exploration when one of the remaining options was chosen. Pairwise post-hoc contrasts of fixed effects were performed using MATLAB’s `coefTest` function.

## 3 RESULTS

### 3.1 Model-agnostic measures (sample 1, de novo dataset, *N* = 75, mixed sex)

Analyses of differences in model-agnostic measures between drug conditions (see section 2.3.1 for details on the model-agnostic trial classification scheme) did not support the hypothesis that L-DOPA would increase response vigour, i.e., reduce RTs, as Bayes factors showed strong evidence for a drug-related increase in RT (BF₀₊ = 22.09). Further, the analyses yielded only moderate evidence supporting our prediction of reduced uncertainty-based exploration (BF₀₊ = 6.28; placebo: *M* = 0.15, L-DOPA: *M* = 0.14). For total points, random exploration, and optimal choices, the two-sided Bayes’ factors (BF₀₁ = 7.69–7.74) indicate moderate evidence in favour of the null hypothesis. Bayes’ factors provided moderate evidence against the hypothesised increase in stay trials (BF₀⁻ = 5.30) and decrease in switch trials (BF₀⁺ = 5.21) under L-DOPA. Assessing the relationship between RTs and choice type, we found strong evidence that mean RTs were faster for directed compared to random exploration trials (BF_10_ = 22.15; *M*_directed_ = 0.45, *M*_random_ = 0.46). Full results are reported in supplementary Table S6 and Figure S7.

### 3.2 Model comparison

The model comparison was conducted on sample 1 data (de novo dataset, *N* = 75, mixed sex). LOO cross-validation scores (Vehtari et al., 2017) for each model and drug condition revealed an overall superior fit of the Bayesian learner models (see Table 3). Here, variants including terms for directed exploration, higher-order perseveration, and total uncertainty (SM_+E +HOP +TU_) outperformed the other models with regard to predictive accuracy in both drug conditions (see Table 3). This model (Bayesian learner with SM_+E +HOP +TU_) was therefore implemented as a combined model across drug conditions, in which the placebo parameters were modelled as the baseline and parameters in the L-DOPA condition as additive drug effects.

### 3.3 Posterior predictive checks

We evaluated model performance of the best-fitting model (Bayesian learner with SM_+E +HOP +TU_) with posterior predictive checks by simulating 4,000 choices per trial and participant from the posterior distribution and comparing them to observed choices in terms of accuracy (predicted-observed match), switch rate, and proportion of optimal choices, separately for each random walk (see Figure 1). Posterior predictive checks showed a good overall correspondence between predicted and observed choices, confirming that the model captured key aspects of participants’ choice behaviour. Notably, for random walk 2, switch rates decreased toward the end of the experiment, while optimal choice rates increased. See supplementary Figure S8 for posterior predictive checks for individual participants across trials.

**Figure 1.**
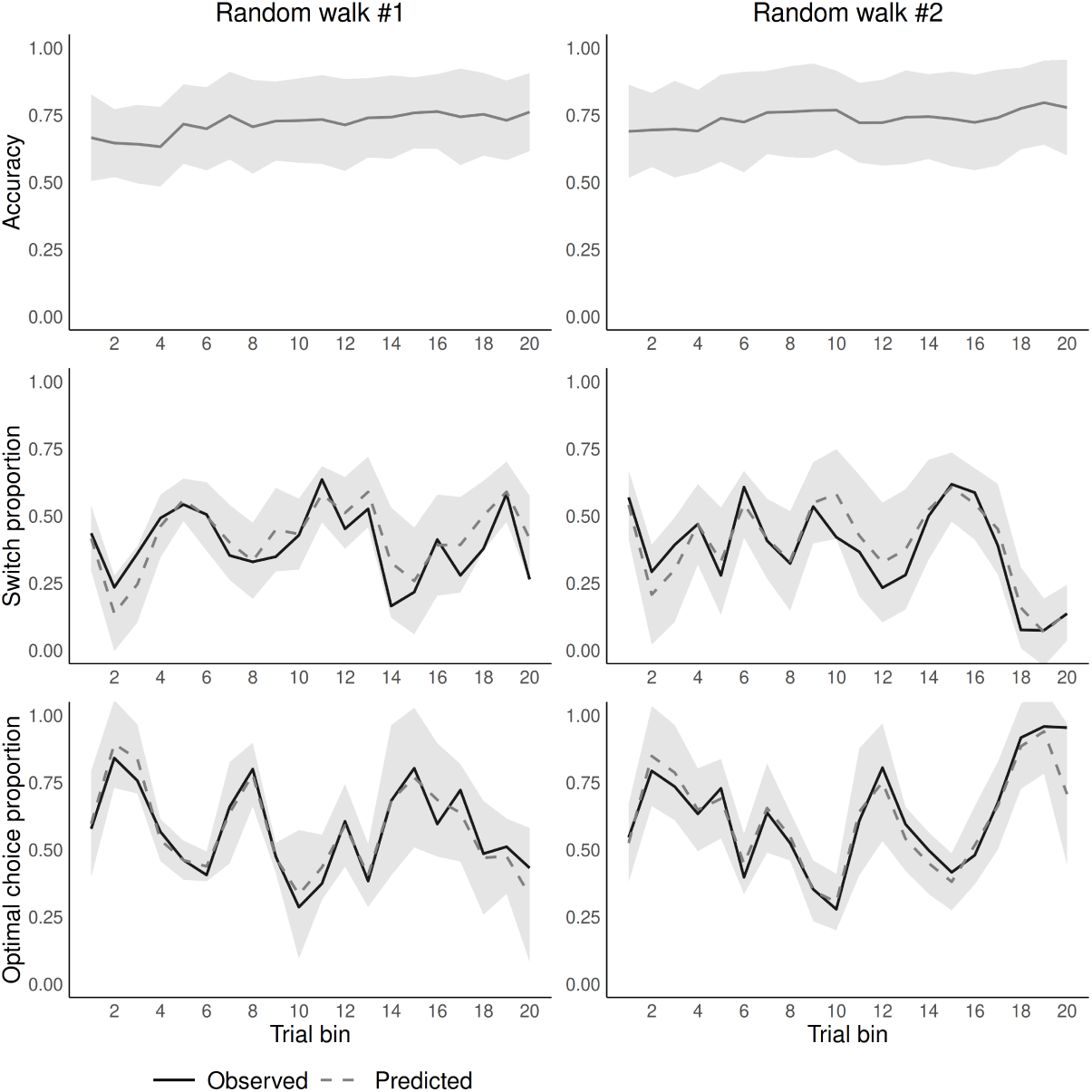
Posterior predictive checks. Comparison of simulated and observed choices, separately for each random walk, averaged across simulations and grouped into 20 bins of 10 trials each (200 trials in total). Top row: Predictive accuracy, i.e., proportion of trials where the model’s choice prediction matched the observed choice; middle row: proportion of switches, i.e., trials where participants chose a different option than on the previous trial; bottom row: Proportion of optimal choices, i.e., selections of the objectively best option. Solid lines: mean observed data across participants; dashed lines: model simulations from the posterior distribution (mean across 4,000 simulations); ribbons: standard deviations of posterior simulations.

### 3.4 Drug effects on model parameters in the SM_+E +HOP +TU_ model

#### 3.4.1 Sample 1 (de novo dataset, *N* = 75, mixed sex)

In the placebo condition, we observed the predicted effects of directed exploration and HOP. Specifically, *φ* was positive, consistent with our prediction that participants would engage in directed exploration. Similarly, *α _HOP_* was positive though below one, reflecting traces of HOP (see Figure 2), i.e., an influence of prior choices beyond the immediately preceding one. Notably, under placebo, the effect of total uncertainty (*γ*) was negative, indicating that participants initially relied more heavily on value estimates when total uncertainty was high and less when total uncertainty was low. Inspecting the posterior distributions of the drug effect parameters, we found no credible effect of L-DOPA on *β* (softmax inverse temperature, i.e., random exploration), *φ* (directed exploration), *ρ* (perseveration strength), *α _HOP_* (habit update) or *γ* (total uncertainty). In all cases, the 95% HDIs of the posterior distributions included zero (see Table 4 and Figure 2). Extending our preregistered analyses, we checked for a potential association between the drug effects and body weight, but observed no credible evidence for body weight effects (see supplementary Figure S9). We further conducted the analysis with the sample split by body weight (median split). Again, in neither BMI group did we observe credible evidence for drug effects, as the 95% HDIs included zero in all cases (see supplementary Figure S10).

**Figure 2.**
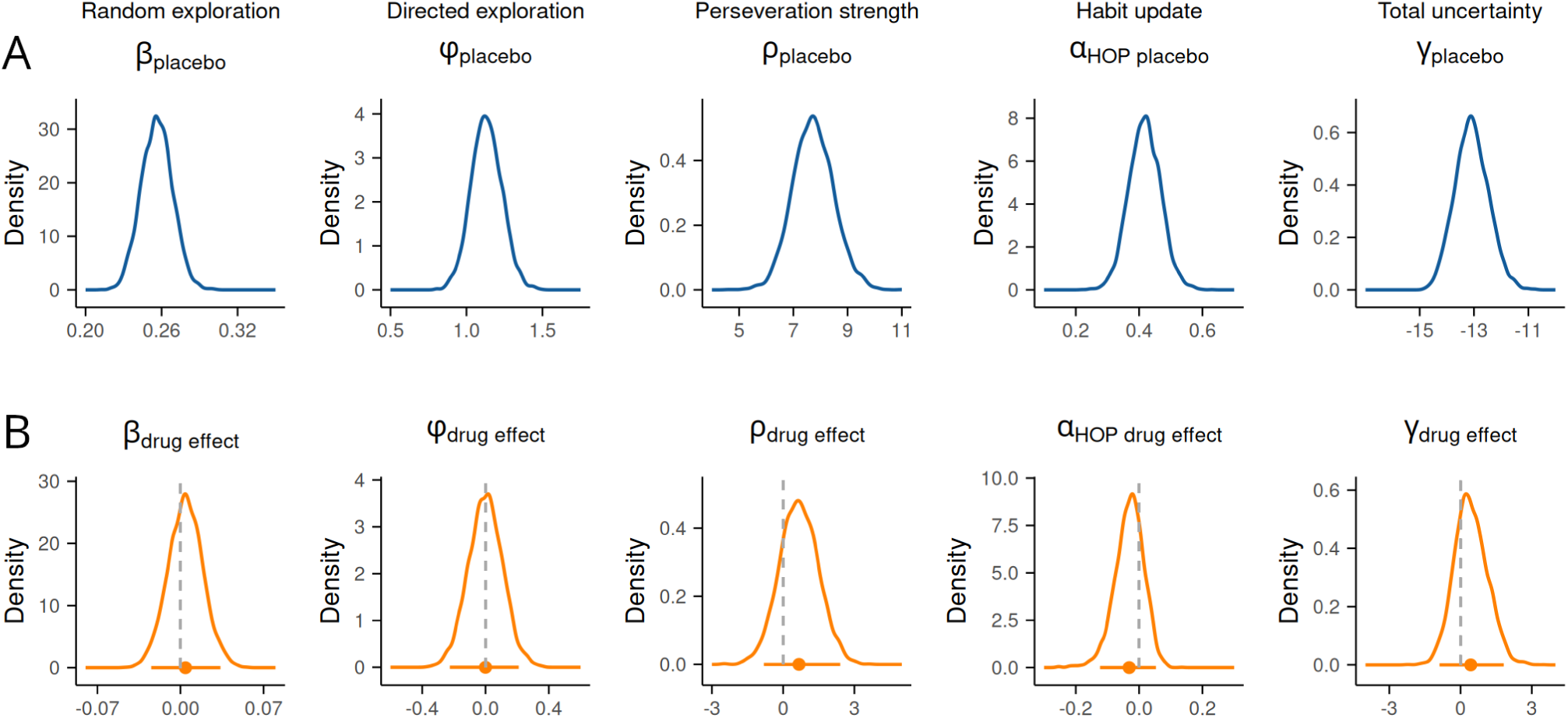
Posterior distributions of the group-level parameter means of sample 1 (*N* = 75, mixed sex) for the best-fitting Bayesian learner model including terms for including terms for random and directed exploration, higher-order perseveration and total uncertainty (SM_+E +HOP +TU_). Panel A (top row, blue distributions): parameters in placebo condition modelled as baseline. Panel B (bottom row, orange distributions): drug effects modelled as additive changes from placebo to L-DOPA. Vertical dashed line: *x*=0. Horizontal solid orange line: 95% highest posterior density interval. *β*: random exploration (softmax inverse temperature); *φ*: directed exploration; *ρ*: perseveration strength; *α _HOP_*: habit update; *γ* : total uncertainty weight.

**Table 4.**
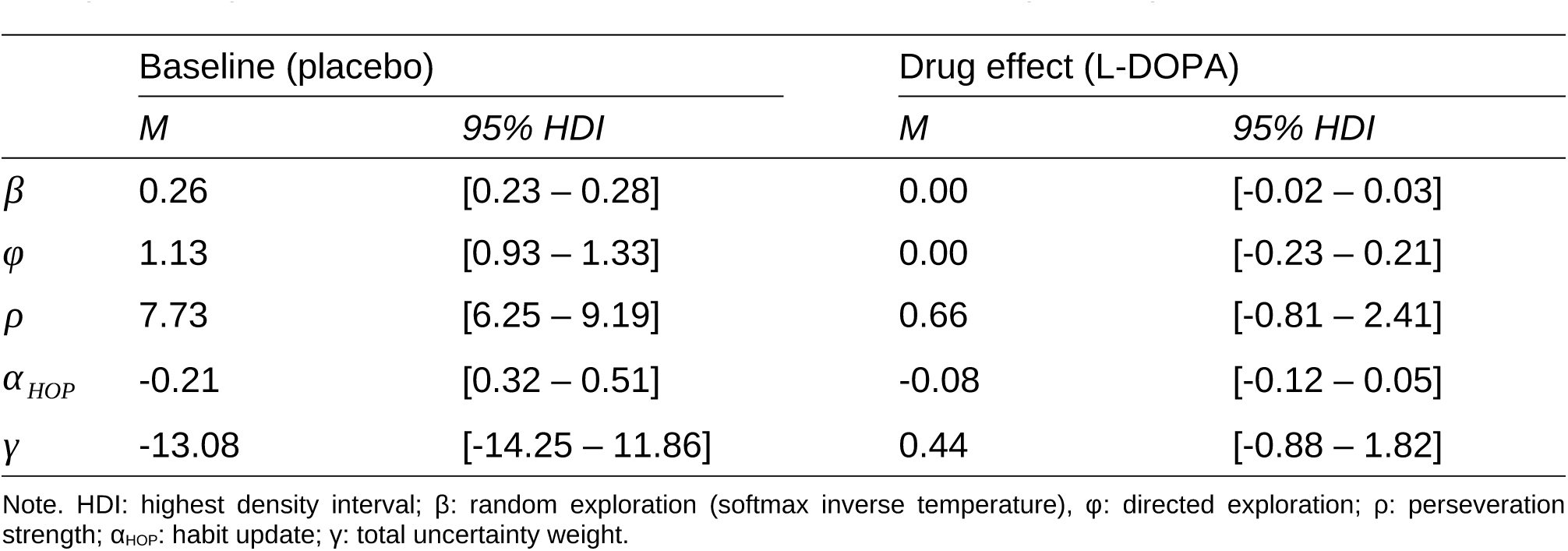
Summary of model parameters (means and 95% HDIs) of the drug-effect Bayesian learner model SM_+E +HOP +TU_. Parameters in the placebo condition were modelled as baseline, while changes from placebo to L-DOPA were modelled as additive drug-effect parameters.

|  | Baseline (placebo) |  | Drug effect (L-DOPA) |  |
| --- | --- | --- | --- | --- |
|  | *M* | 95% *HDI* | *M* | 95% *HDI* |
| $\beta$ | 0.26 | [0.23 – 0.28] | 0.00 | [-0.02 – 0.03] |
| $\phi$ | 1.13 | [0.93 – 1.33] | 0.00 | [-0.23 – 0.21] |
| $\rho$ | 7.73 | [6.25 – 9.19] | 0.66 | [-0.81 – 2.41] |
| $\alpha_{HOP}$ | -0.21 | [0.32 – 0.51] | -0.08 | [-0.12 – 0.05] |
| $\gamma$ | -13.08 | [-14.25 – 11.86] | 0.44 | [-0.88 – 1.82] |

Note. HDI: highest density interval; $\beta$ : random exploration (softmax inverse temperature), $\phi$ : directed exploration; $\rho$ : perseveration strength; $\alpha_{HOP}$ : habit update; $\gamma$ : total uncertainty weight.

#### 3.4.2 Sample 2 (pooled sample, *N* = 74 men)

Next, we pooled the male participants from the de novo dataset with the *n* = 31 participants (all male) from Chakroun et al. (2020) and again fitted the SM_+E +HOP +TU_ model. Posterior distributions of the drug effect parameters and their 95% HDIs indicated a credible positive drug effect on *ρ* (perseveration strength) and a negative drug effect on *α _HOP_* (habit update), (see Figure 3), such that L-DOPA increased the tendency to repeat previous choices and reduced the decay of their influence over time, thereby extending the window of choice history integration on current decisions. Further, we found a credible positive drug effect on *γ* (total uncertainty), such that L-DOPA increased uncertainty-dependent value weighting. Notably, the baseline (placebo) *γ* parameter was negative, indicating that participants initially relied more heavily on value estimates when total uncertainty was high and less when uncertainty was low.

**Figure 3.**
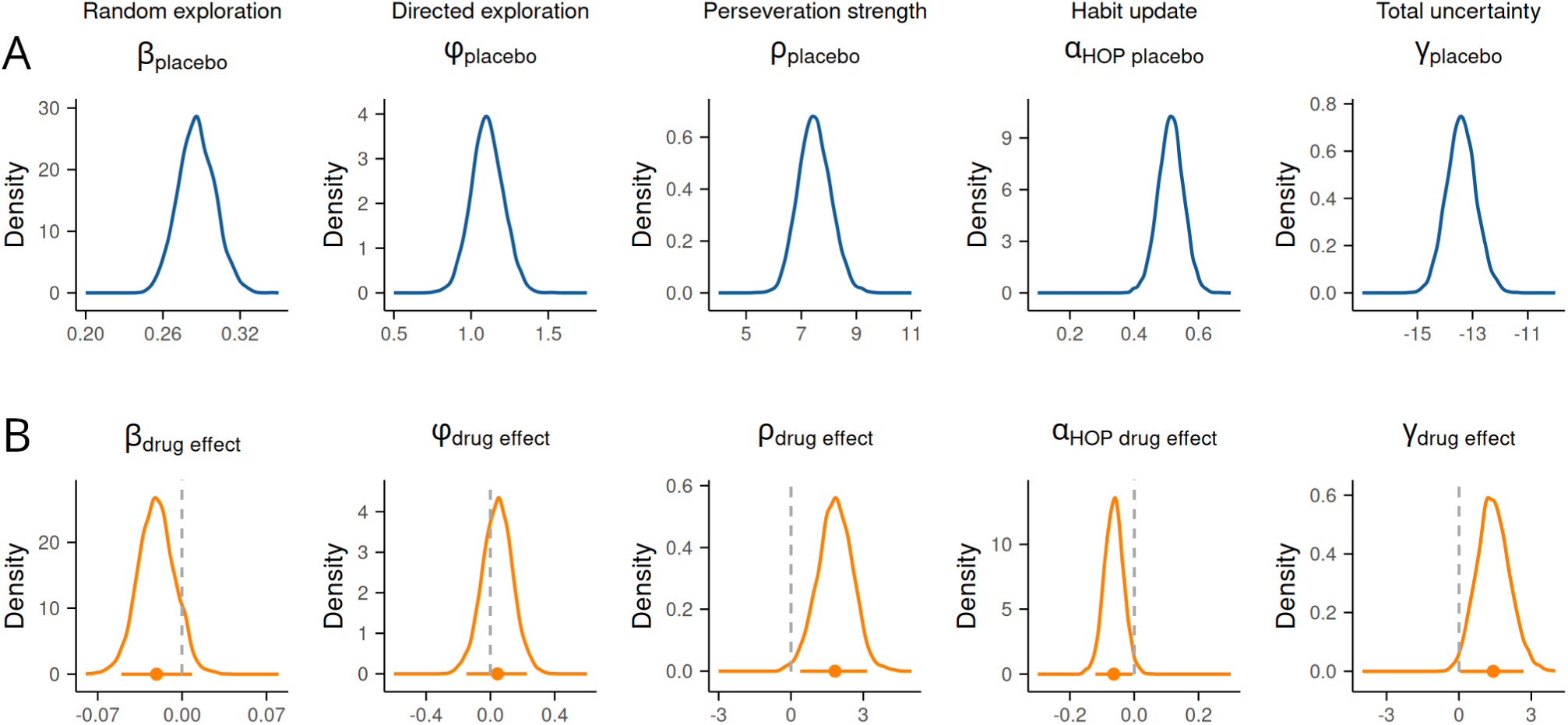
Posterior distributions of the group-level parameter means of sample 2 (pooled male-only sample, *N* = 74) for the best-fitting Bayesian learner model including terms for random and directed exploration, higher-order perseveration and total uncertainty (SM_+E +HOP +TU_). Data for the male group were pooled with participants from Chakroun et al. (2020), who used the same task in a male-only sample. Panel A (top row, blue distributions): parameters in placebo condition modelled as baseline. Panel B (bottom row, orange distributions): drug effects modelled as additive changes from placebo to L-DOPA. Vertical dashed line: *x*=0. Horizontal solid orange line: 95% highest posterior density interval. *β*: random exploration (softmax inverse temperature); *φ*: directed exploration; *ρ*: perseveration strength; *α _HOP_*: habit update; *γ*: total uncertainty weight.

The positive L-DOPA effect on *γ* reflects that L-DOPA increased uncertainty-dependent value weighting, enabling participants to rely more on value estimates when uncertainty about options was low, and less when uncertainty was high. In the subsample of female participants from the de novo dataset (*n* = 32) we found no credible drug effects for any of the parameters (see supplementary Figure S11). The difference between male (sample 2) and female (sample 1) posterior distributions for each model parameter is shown in Supplementary Figure S12. There was no credible evidence for sex differences in drug effects for any parameter. However, for *β*, the overlap of the HDI with zero was marginal (HDI = [-0.001, 0.087], suggesting higher random exploration in female vs. male participants.

### 3.5 Drug effects and interindividual differences (sample 1, de novo dataset, *N* = 75, mixed sex)

We assessed the relationship between drug effects and DA proxy measures (BIS-15 scores, sEBR, WM capacity) using Bayesian regression, testing both linear and non-linear (quadratic) effects by regressing the drug effect parameters on the (squared) DA proxies. The analysis revealed a credible positive effect of WM capacity on *β* (softmax inverse temperature, i.e. random exploration), with both the linear and quadratic coefficients positive, as indicated by 95% HDIs not including zero (see Figure 4A). We found no credible evidence that any of the other DA effect parameters were associated with the DA proxy measures, neither in a linear nor quadratic fashion, as all 95% HDIs included zero (see Figure 4B-E). The relationships between DA proxies and drug-effect parameters are shown in Figure 5, including individual data points and fitted linear and quadratic curves. To assess the robustness of the original Bayesian regression results relating drug effects to dopamine-proxy measures and WM capacity, we repeated the analysis after excluding two participants whose WM capacity scores fell more than 2 SDs below the sample mean. The overall pattern of results remained consistent with the full dataset. While the quadratic term for WM capacity was not credibly different from zero, the linear effect remained credibly positive (see Supplementary Figure S13). Since we found drug effects in male but not female participants (see section 3.4), and observed that drug effects were mediated by WM capacity, we tested for sex differences in WM capacity. However, for all measures, Bayes’ factors (calculated in JASP using Bayesian Mann-Whitney U tests) favored the null hypothesis, indicating no evidence for sex differences, neither for overall WM capacity, operationalised as the first principal component across all three WM tasks (BF₁₀ = 0.32, W = 580), nor for any of the individual WM components, i.e., digit span forward (BF₁₀ = 0.26, *W* = 635), digit span backward (BF₁₀ = 0.52, *W* = 521), listening span (BF₁₀ = 0.26, *W* = 608), or operation span (BF₁₀ = 0.42, *W* = 570).

**Figure 4.**
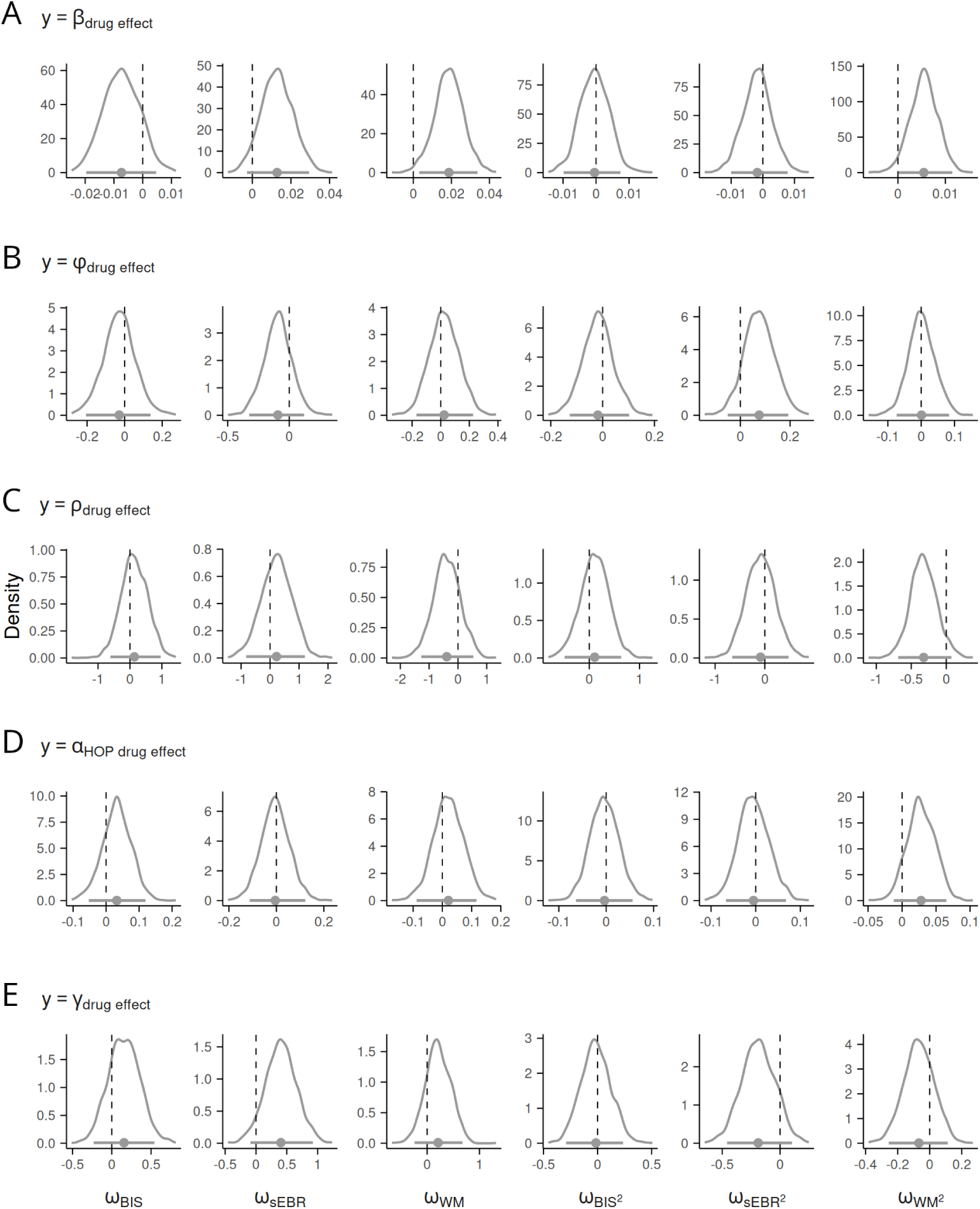
Posterior distributions of the linear (column 1 to 3) and quadratic (column 4 to 6) regression coefficients (*ω*), regressing the drug effect parameters of the Bayesian learner model with SM_+E +HOP +TU_ onto the (squared) putative DA proxy measures impulsivity (BIS-15 score), sEBR and WM capacity. The posterior distribution of the intercept is omitted. Vertical dashed line: *x*=0. Horizontal solid line: 95% highest posterior density interval. *s*: drug effects modelled as additive changes from placebo to L-DOPA. *β*: random exploration (softmax inverse temperature); *φ*: directed exploration; *ρ*: perseveration strength; *α _HOP_*: habit update; *γ*: total uncertainty weight; BIS-15: scores for the Barratt Impulsiveness Scale (short form); sEBR: spontaneous eye blink rate, WM: working memory capacity.

**Figure 5.**
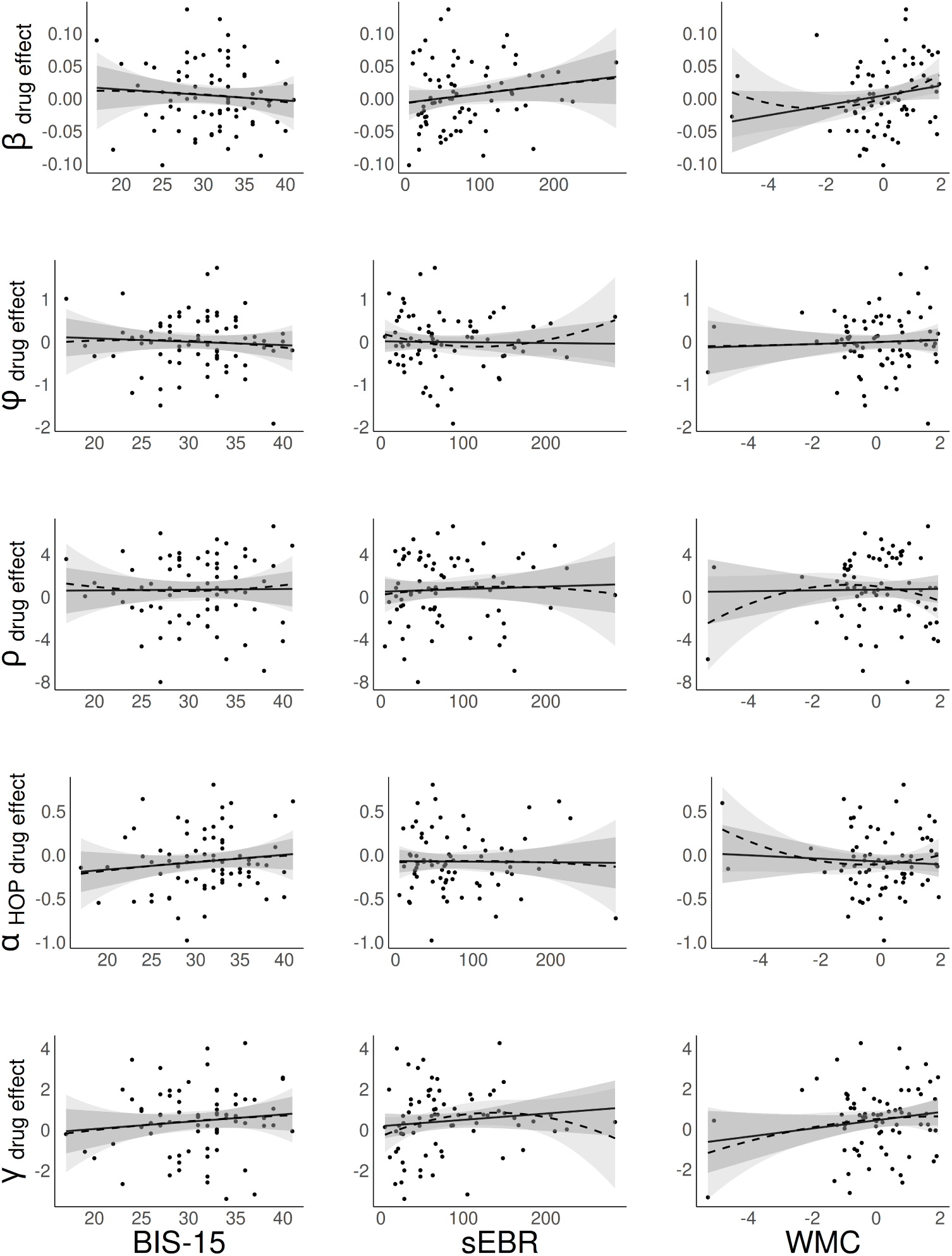
Relationship between the putative DA proxy measures impulsivity (BIS-15 scores), spontaneous eye blink rate (sEBR) and working memory capacity (WMC) and the drug effects (means of single-subject posterior distributions for the best-fitting Bayesian learner model including terms for random and uncertainty-based exploration, first and higher-order perseveration and total uncertainty (SM_+E +HOP +TU_). Points: single-subject values; solid black line: linear fit; dashed line: quadratic fit; grey ribbons: standard error. *β*: softmax inverse temperature (random exploration); *φ*: directed exploration; *ρ*: perseveration strength; *α _HOP_*: habit update; *γ*: total uncertainty weight.

### 3.6 Visual fixation and pupil dilation

A GLMM assessing the relationship between the number of fixation shifts before making a choice and uncertainty revealed that uncertainty of the chosen option was associated with an increase in fixation shifts (β = 0.071, *SE* = 0.013, *p* < .001), while total uncertainty was associated with fewer fixation shifts (β = −0.032, *SE* = 0.019, *p* < .001). These effects were not significantly associated with drug condition (all main effects and interactions *p* > .05; see Supplementary Table S14 for full statistical reporting of effects on fixation shifts). We observed a significant association between the last fixation target and the choice made (χ² = 53,454, *p* < .001), confirming that prior fixations are predictive of choices. On most trials, participants fixated an option once and chose it subsequently (94%). In 5% of trials, participants shifted fixation once, i.e., fixated two different options before making a choice. Only a small percentage of trials involved more frequent fixation shifts between the options, with 0.6% involving two shifts, and 0.11% involving three or more shifts.

Pupil dilation (AUC, see methods section) was analysed using GLMMs for each trial phase with effects of drug, trial number, and decision variables. As predicted, L-DOPA increased pupil dilation relative to placebo, but this effect was observed only during the active phases of the task (*p* < .01 for the pre-choice and feedback phase; *p* = .108 for the inter-trial-interval; see supplementary Table S15 for full reporting of GLME coefficients and statistical results for each task phase). We found a significant negative effect of trial number during the pre-choice phase and inter-trial-interval (*p* < .001), reflecting that pupil diameter decreased progressively across trials. No significant interactions were observed between drug and trial (*p* > .05 for all trial phases). Higher expected value of the chosen option was associated with reduced pupil dilation (*p* < .01 for all trial phases, an effect for which we had no specific hypotheses). Also, higher uncertainty of the chosen option and total uncertainty was associated with reduced pupil dilation across all task phases (*p* < .01 for all phases). As predicted, pupil dilation scaled with prediction error (both signed and unsigned) during feedback, and this effect extended into the inter-trial interval (*p* < .05 for signed prediction error; *p* < .001 for unsigned prediction error). Further, consistent with our prediction, pupil diameter varied with choice type (note that we did not make phase-specific hypotheses for choice type modulation; see section 2.3.8 for details on the model-based trial classification scheme). As hypothesised, pupil dilation was larger for random exploration compared to exploitation before making a choice (*p* < .001) and during the inter-trial-interval (*p* = .001). In contrast, no significant differences were found for directed exploration compared to exploitation (*p* > .05 for all phases). Differences between directed and random exploration were not significant after correcting for multiple comparisons (*p* > 0.017 for all posthoc tests). See Figure 6 for pupil responses under placebo vs. L-DOPA, and for the relationships between GLME-estimated pupil responses and decision variables for each trial phase.

**Figure 6.**
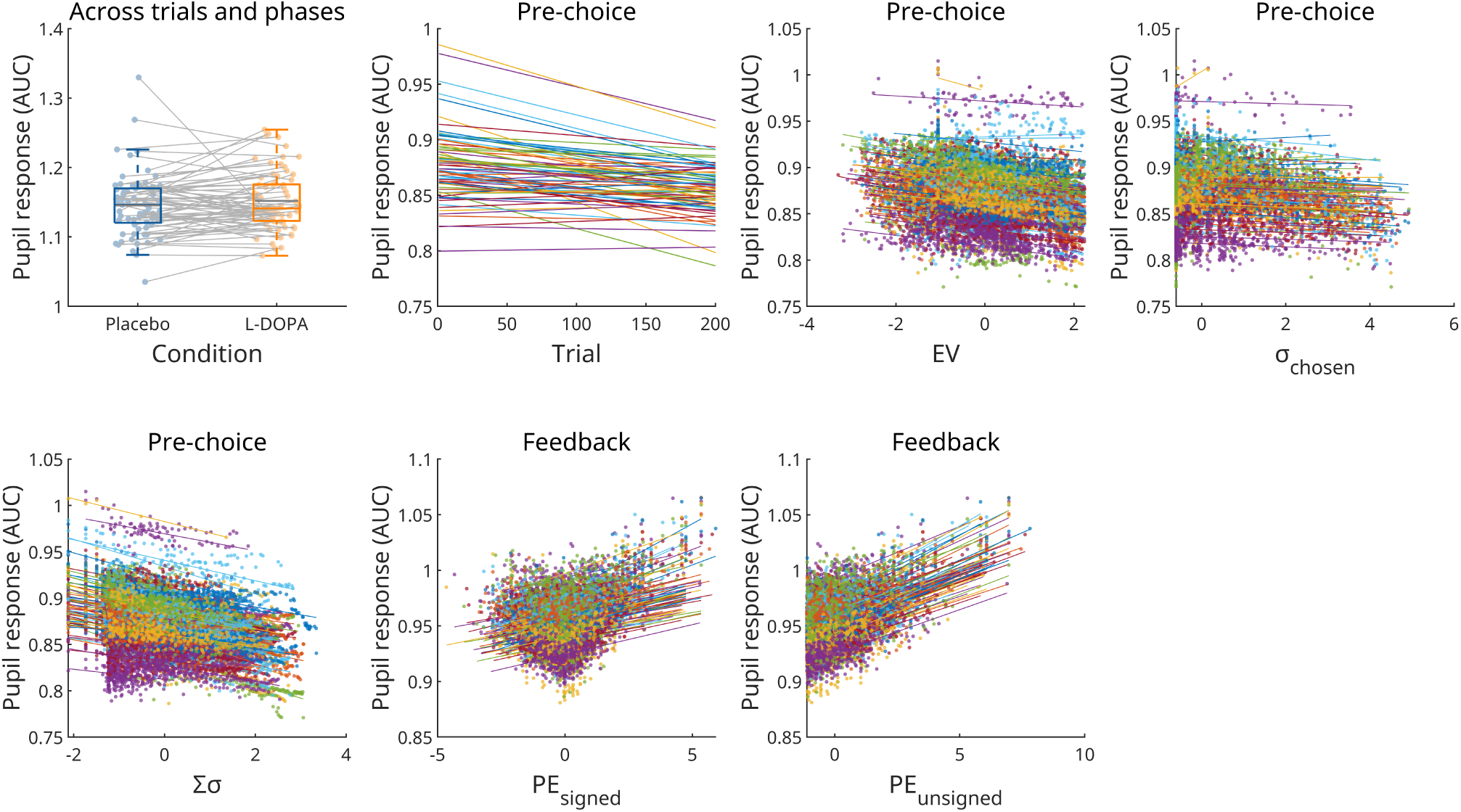
Panel 1: participant-wise mean pupil dilation (area under the curve, AUC) averaged across all task phases for the placebo and L-DOPA condition (sample 1 data). Solid lines connect data points from the same participant across conditions. Boxes: interquartile range (IQR); whiskers 1.5 × IQR; horizontal coloured lines: median; horizontal grey line: mean. Panel 2: Subject-level predicted trajectories of pupil dilation across trials during the pre-choice phase, illustrating within-and between-participant variability in baseline and slope estimates. Panel 3–7: generalised linear mixed-effects (GLME) estimates of pupil dilation (area under the curve, AUC) from sample 1 plotted against trial-wise decision variables during the pre-choice and feedback phase. Each colour represents an individual participant’s predicted effect, with linear fits overlaid. All continuous predictors (i.e., uncertainty measures and decision variables) were standardised. AUC: area under the curve, EV: expected value; *σ_chosen_*: uncertainty of chosen option; *Σσ* : total uncertainty; PE: prediction error.

## 4 DISCUSSION

We investigated the role of DA in modulating exploration-exploitation behaviour during RL. Although pharmacological evidence consistently linked DA to exploration, inconsistencies in the directionality of the effects suggest that baseline DA function may modulate drug effects. We therefore tested the effects of increasing DA signalling using L-DOPA in a randomised, double-blind, placebo-controlled study (*N* = 75) and examined the moderating influence of putative baseline DA markers, including sEBR, impulsivity, and WM capacity. In addition, we assessed visual fixation patterns and pupil dilation as physiological correlates of decision-making.

### Model comparison

Our model expanded on previous modelling approaches for restless bandit tasks (Chakroun et al., 2020; Wiehler et al., 2021) in two ways. First, we included a total uncertainty term (Gershman, 2018; Lau & Glimcher, 2005), i.e., uncertainty across all available options. Second, based on several recent suggestions and model comparison work (Brands et al., 2025), we included a higher-order perseveration (HOP) term to model choice repetition effects beyond trial t-1 (Miller et al., 2019). Model comparison using LOO cross-validation (Vehtari et al., 2017) showed that an extension of a Bayesian learner model (Chakroun et al., 2020; Wiehler et al., 2021) with higher-order perseveration and a total uncertainty term (SM_+E +HOP +TU_) provided the best fit under both drug conditions. Posterior predictive checks confirmed that the model accurately reproduced the observed time courses of accuracy, switching behaviour and optimal choices across trials.

### DA effects on exploration and interindividual differences

In the full model across drug conditions, we observed no credible evidence for L-DOPA effects on model parameters in our de novo mixed-sex dataset. This contrasts with our earlier study (Chakroun et al., 2020), where L-DOPA reduced directed exploration in *n* = 31 healthy men. Since we used a fixed dose, differences in effective dosage may partly explain this. However, control analyses taking into account body weight also did not reveal credible effects (see supplementary Figure S9). We pooled male participants from sample 1 and participants from Chakroun et al. (2020) to examine potential sex-specific effects. Placebo parameter estimates were similar between sample 1 and sample 2, indicating some degree of consistency across datasets. Our final model, which extended Chakroun et al. (2020) by including higher-order perseveration and total uncertainty terms, revealed notable differences in drug effects. In the new model, we did not observe the predicted effect of L-DOPA on directed exploration. Instead, in sample 2 (pooled male-only sample), L-DOPA increased perseveration and reduced habit updating, indicating a stronger tendency to repeat past choices and a slower decay of their influence over time (increased temporal integration). One possibility is that these mechanisms trade off with directed exploration, which may in part explain the discrepancy with Chakroun et al. (2020). Whereas that study found reduced directed exploration, our results suggest increased perseveration under L-DOPA in men. However, both patterns reflect reduced flexibility, consistent with a shift towards reduced exploration. DA is implicated in cognitive flexibility in healthy and clinical populations (Klanker et al., 2013) and, in line with this, DA agonists increased perseveration in Parkinsonian rats (Faure et al., 2010).

In men, L-DOPA also influenced uncertainty-dependent weighting of value estimates, captured by the total uncertainty weight (*γ*). Under placebo, participants exhibited negative γ values, indicating they relied more heavily on value estimates when total uncertainty across options was high, and reduced their reliance on value estimates when uncertainty was low. This represents a suboptimal strategy where individuals do not calibrate their exploitation behaviour to their uncertainty, i.e., under-weighting their most reliable value estimates when they should have trusted them most. L-DOPA increased γ, promoting a pattern of stronger uncertainty-calibrated decision-making. Participants relied more strongly on value estimates when uncertainty was low (confident exploitation) and reduced value-based decision-making when uncertainty was high, allowing other components to drive behaviour more strongly. This may reflect a more optimal regulation of behaviour contingent on uncertainty (although overall performance scores were similar between drug conditions). These findings suggest that DAergic enhancement improved metacognitive regulation of decision strategies, enabling participants to better use uncertainty signals to balance exploration and exploitation. Importantly, this increase in uncertainty-calibrated decision-making occurred alongside L-DOPA’s enhancement of perseveration. While these effects may seem contradictory, they likely reflect DAergic modulation of distinct but complementary decision-making processes operating at different timescales. The effect on total uncertainty represents enhanced within-trial metacognitive control, i.e., better moment-to-moment calibration of exploitation to uncertainty. Meanwhile, increased perseveration reflects stronger across-trial commitment to recently chosen policies. Together, these findings suggest that L-DOPA may have enhanced both the refinement of immediate decision strategies and the stability of longer-term behavioural patterns, potentially reflecting DAergic effects on prefrontal metacognitive processes (Ershadmanesh et al., 2025; Joensson et al., 2015; Qiu et al., 2018) and striatal habit formation (Greenstreet et al., 2025; van Elzelingen et al., 2022), respectively.

We also tested the degree to which cognitive processes putatively linked to baseline DA influenced L-DOPA effects, as predicted by the inverted-U hypothesis of DA function (Cools & D’Esposito, 2011). Here, WM capacity, but not sEBR and impulsivity, was credibly associated with drug effects. Notably, this null finding for sEBR is consistent with prior work reporting no association between sEBR and the effects of a DA agonist (bromocriptine; Dang et al., 2017). Specifically, although our model revealed no overall drug effect on random exploration (softmax inverse temperature), WM capacity positively modulated this effect (de novo mixed-sex sample).

The linear term remained robust after excluding outliers, supporting the notion that the effect of L-DOPA may interact with pre-existing cognitive capacities to shape choice behaviour. L-DOPA may further stabilise value representations, e.g., in prefrontal regions (Durstewitz et al., 2000), leading to more deterministic choices. This dovetails with previous reports of more pronounced L-DOPA effects on (goal-directed) decision-making in individuals with high WM capacity (Kroemer et al., 2019). This pattern also fits with our recent recurrent neural network (RNN) modelling results, showing increased directed exploration and attenuated random exploration in high vs. low-capacity RNNs, suggesting that greater representational capacity promotes more goal-directed and less stochastic choice behaviour (Flimm et al., 2025). Further, high-WM individuals may differ in DA receptor availability (Rieckmann et al., 2011), which may lead to a differential sensitivity to pharmacological interventions. Note, however, that the effect size of the association between WM capacity and random exploration was small. Also, in our previous report based on sample 1 temporal discounting data (Smith et al., 2025), we did not observe credible evidence for an association of WM capacity with L-DOPA effects on model parameters. This suggests that this link may be task-specific, small and potentially not robust.

In summary, L-DOPA increased sensitivity to uncertainty and perseveration, and reduced habit updating (increased temporal integration) in the male-only sample (pooled with Chakroun et al., 2020), suggesting a potential influence of sex on these processes. This aligns with previous research showing that noradrenergic effects on exploration are sex-dependent in mice (Chen et al., 2024), and that DAergic signaling and drug responsiveness also vary by sex, partly due to hormonal modulation (Sinclair et al., 2014). Of note, sex differences in dopaminergic drug effects have also been documented in the clinical literature, where dopaminergic treatment is associated with sex-dependent prognosis of impulse control disorders in Parkinson’s disease (Joutsa et al., 2012). While we restricted female participants to those using hormonal contraception to reduce variability, this does not eliminate hormonal influences. This variability may have masked drug effects, as DA signalling can fluctuate with the menstrual cycle (Czoty et al., 2009; Dreher et al., 2007). Since credible drug effects emerged in the male sample only, we also tested for sex differences in WM capacity (Harness et al., 2008; Voyer et al., 2017; Zilles et al., 2016). This revealed no credible sex differences, neither for the composite WM measure (PC1) nor the individual measures. Nonetheless, men and women may partly rely on different cognitive strategies during RL (Chen et al., 2021; Evans & Hampson, 2015), which could differentially modulate L-DOPA’s effect. Sex-related variation in pharmacokinetics and metabolism (Soldin & Mattison, 2009; Walker et al., 2006) and DA synthesis capacity (Laakso et al., 2002; Munro et al., 2006; Pohjalainen et al., 1998) may also contribute to sex-specific drug responses. The smaller female subgroup in sample 1 further decreased power to detect subtle effects. While a stratified analysis revealed credible drug effects in men, but not in women, when directly comparing the posterior distributions of drug effects between men and women, the differences were not credibly different from zero. This implies the effect is present in men, but does not credibly differ from the effect in women. Together, this highlights the complexity of sex-dependent DAergic modulation and underscore the need for further research into sex-specific mechanisms in pharmacological interventions.

### Fixation and pupil dilation correlates of exploration

We also examined how decision variables and L-DOPA influenced visual fixation patterns and pupil dilation, to gain further insight into the underlying mechanisms. Analysis of fixation shifts (i.e., the number of gaze shifts between options) revealed a negative association between total uncertainty and fixation shifts. Total uncertainty is directly linked to the length of the current exploitation streak. This effect therefore likely reflects reduced visual exploration during longer periods of exploiting a specific option. In contrast, fixation shifts increased with increasing uncertainty of the chosen option, reflecting visual exploration during exploratory choices. L-DOPA did not credibly influence uncertainty-related visual sampling. Final fixations reliably predicted choice, consistent with prior work (Isham & Geng, 2013) and in line with the broader view that visual attention is a central driver of decision-making (Krajbich & Rangel, 2011; Moerel et al., 2024; Orquin & Loose, 2013; Summerfield & Egner, 2014).

As predicted, L-DOPA increased pupil dilation compared to placebo, consistent with prior results (Bartošová et al., 2018; Spiers & Calne, 1969), and providing physiological evidence for an overall drug response in sample 1 participants. In line with our hypothesis, pupil dilation scaled with prediction error during feedback (Preuschoff et al., 2011), increasing with both signed and unsigned prediction error, but more strongly with the latter. This indicates that surprise dominated the pupil response, while value learning also played a role. In line with our hypothesis, and consistent with previous reports (Gilzenrat et al., 2010; Jepma & Nieuwenhuis, 2011), pupil dilation varied with choice type such that random exploration was associated with greater dilation compared to exploitation. However, this was not the case for directed exploration vs. exploitation nor directed vs. random exploration. Although directed exploration may involve greater deliberate processing compared to random exploration (Wilson et al., 2021), pupil responses were not sensitive to this effect. Further, pupil dilation was negatively associated with expected value, contrasting with previous findings (Van Slooten et al. 2018). A possible explanation is that our four-armed restless bandit task differs fundamentally from the probabilistic selection task used by Van Slooten et al. (2018), in which stimulus-reward contingencies are fixed and learned gradually. In our task, because the reward contingencies drift, choosing a low-expected value option may often be a strategic choice to sample an uncertain option. Therefore, larger pupil responses may reflect the informational value or cognitive effort required to deviate from the exploit-state, rather than the low value itself.

Together, these findings indicate that pupil dilation reflects both cognitive and pharmacological effects during exploration-exploitation decisions. Phasic changes tracked expected value, uncertainty, prediction error, and choice type, while L-DOPA exerted a tonic dilating effect during active trial phases, i.e., not significantly during the inter-trial interval. This absence may reflect reduced engagement and arousal, limiting both baseline variability and sensitivity to DAergic effects. Finally, drug, cognitive variables, and choice type effects on pupil dilation were statistically credible, with the effect of the drug on pupil dilation comparatively stronger. However, all effects were relatively small in magnitude. Thus, while pupil dilation reflects cognitive and pharmacological effects, it may be constrained in complex task settings with overlapping influences. While pupil dilation provides a coarse index of cognitive processing, fixation patterns may offer a more detailed view of how participants sample information under uncertainty.

### Limitations

This study has several limitations. First, as participants initially completed a temporal discounting task (of approx. 12 minutes duration), drug timing may have influenced the results due to the drug’s half-life. Following oral administration, L-DOPA reaches peak plasma concentrations within 30–60 minutes and maintains a half-life of approximately 90 minutes (Hauser, 2009; Keller et al., 2011). The fact that we observed a drug effect on pupil responses suggests that pharmacological activity was still present. Second, we did not directly measure DA function. The study was designed to examine the effects of a DA challenge on RL, and how this might interact with interindividual differences in putative proxy measures of baseline DA. However, a recent PET study found no link between striatal DA synthesis capacity and WM capacity, sEBR, or impulsivity (van den Bosch et al., 2023). Our findings further strengthen the notion that these measures might reflect cognitive and/or physiological variation unrelated to baseline DA. However, in our study, WM capacity was associated with L-DOPA’s effects on random exploration. In light of the absence of a link between WM capacity and DA synthesis capacity (van den Bosch et al., 2023), this may be due to other factors influencing DAergic drug responses. For example, this could be due to other aspects of DA function beyond synthesis capacity, such as DA receptor sensitivity and DA release. Furthermore, L-DOPA’s effects on exploration could be mediated by DA’s role in reward processing and decision-making (Rutledge et al., 2015; Webber et al., 2021), where WM capacity may contribute to the ability to integrate and utilise reward-related information more efficiently. Thus, even without a direct link between WM capacity and DA synthesis capacity, cognitive factors may explain why WM capacity modulated the effect of L-DOPA on random exploration.

With regard to impulsivity, we may not have detected a relationship with drug effects due to low variability in our sample. Given that we included healthy controls, we may not have captured data from individuals with high impulsivity. Future studies may thus re-examine the relationship in individuals with more extreme or clinically relevant levels. Likewise, generalisability is be limited due to our sample being predominantly WEIRD (Western, Educated, Industrialised, Rich, and Democratic; Henrich et al., 2010).

Individual differences in exploration-exploitation and responsiveness to DAergic drugs may arise from genetic variation in the catechol-O-methyltransferase (COMT) gene, which affects catecholamine degradation (Gershman & Tzovaras, 2018), which was not assessed in the present study. Also, we did not consider effects of menstrual cycle. Since the menstrual cycle may impact on DA D2 receptor availability in monkeys (Czoty et al., 2009) and modulate reward sensitivity in naturally cycling women (Dreher et al., 2007), it may impact on the exploration-exploitation trade-off. To reduce hormonal variability, we included women using hormonal contraception and did not test participants during the pill break. Splitting the female sample by cycle phase was unfeasible due to small subgroup sizes. Further research would help to better understand the influence of hormonal variation. Nonetheless, our study remains a comparatively high-powered investigation with one of the largest samples in comparable pharmacological research.

### Conclusions

This study examined DA effects on exploration-exploitation behaviour during RL. Extending previous modelling approaches, we found that the extension of a Bayesian learner model (Chakroun et al., 2020; Wiehler et al., 2021) with higher-order perseveration (Brands et al., 2025; Miller et al., 2019) and total uncertainty terms (Gershman, 2018) substantially improved model fit. We found no overall drug effects on model parameters in a comparatively large mixed-sex sample (*N* = 75), contrasting with previous findings in a smaller (*N* = 31) male-only sample (Chakroun et al., 2020), and suggesting that DAergic influences may vary by individual factors such as sex. Pooling our male data with Chakroun et al. (2020) revealed that L-DOPA increased perseveration and sensitivity to uncertainty, effects not seen in women (stratified analysis of female participants from sample 1). Participants with higher WM capacity showed a stronger effect of L-DOPA on random exploration, implying that (putative prefrontal) DAergic mechanisms may interact with cognitive capacity in shaping decision-making. Furthermore, L-DOPA increased tonic pupil dilation, while phasic pupil responses reflected prediction errors and choice type. Visual fixation patterns reflected uncertainty-driven sampling, but were unaffected by L-DOPA. Taken together, our findings suggest that DA effects on exploration may depend on WM capacity and sex rather than exerting uniform effects across individuals.

## CODE AND DATA AVAILABILITY

All model code is openly available on the Open Science Framework at https://osf.io/tvxgc. The data cannot be shared publicly, since the participants did not provide consent to public data access. The data are available from RADAR at https://www.radar-service.eu/radar/en/dataset/dxb4zfu6u8mx7u98 (DOI: 10.577 43/dxb4zfu6u8mx7u98), whereby access is granted to researchers for scientific purposes only. The fitted models are openly available at https://zenodo.org/records/15221323.

## AUTHOR CONTRIBUTIONS

**Elke Smith:** Conceptualisation, methodology, software, formal analysis, investigation, data curation, visualisation, project administration, writing - original draft, writing - review & editing

**Hendrik Theis:** Resources, investigation, writing - review & editing

**Thilo Van Eimeren:** Conceptualisation, resources, investigation, writing - review & editing

**Kilian Knauth:** Investigation, data curation, writing - review & editing

**Deniz Tuzsus:** Investigation, writing - review & editing

**Angela Brands:** Methodology, writing - review & editing

**David Mathar:** Methodology, writing - review & editing

**Jan Peters:** Conceptualisation, resources, methodology, project administration, supervision, funding acquisition, writing - review & editing

## ACKNOWLEDGEMENTS

The authors thank Lea Kemalides, Hannah Hacker and Emily Burlon for helping with study organisation and data collection.

