## Supplementary Material for "Individual Differences in Dopaminergic Modulation of Exploration-Exploitation Behaviour"

### S1 Text Principal component analysis (PCA) across WM tasks for sample #1 (*N* = 75)

Since data for the BIS-15 were missing for one participant due to technical issues, the principal component analyses and, based on that, the subsequent analysis of the relationship between the DA proxies (BIS-15 scores, sEBR and WM capacity) and drug effects, is based on data from *n* = 74 participants only. We operationalised WM capacity as the first principal component (PC) derived from three working memory tasks (forward and backward digit span, listening span, and operation span). PCA analysis was conducted using the pca function in MATLAB (version 2022a; The MathWorks, Inc.). All scores were z-scored prior to the analysis. The correlations between the WM scores are shown in Table S2, the cumulative variance explained and the factor loadings are presented in Figure S3. The first PC accounted for 43.3% of the total variance.

S2 Table. Correlations between the WM scores.

| | DS bw max | LS | OS abs |
|---|---|---|---|
| DS fw max | $r = 0.46$, $p < 0.001$ | $r = 0.32$, $p = 0.005$ | $r = 0.22$, $p = 0.064$ |
| DS bw max | | $r = 0.19$, $p = 0.106$ | $r = 0.18$, $p = 0.133$ |
| LS | | | $r = -0.05$, $p = 0.683$ |

Note. WM: working memory; DS: digit span; bw: backward maximum; LS: listening span; OS: operation span; abs: absolute score; fw: forward maximum.

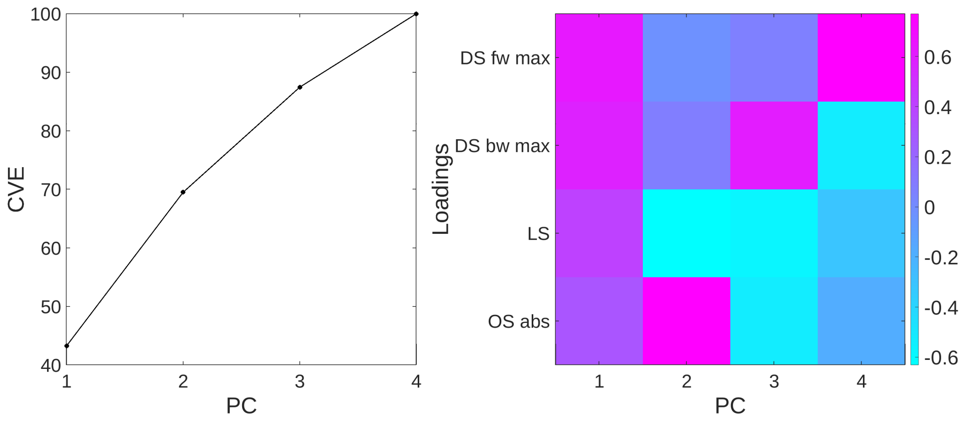


S3 Figure. Results for the PCA across three WM tasks (digit span [DS], listening span [LS], and operation span [OS]). Left panel: cumulative variance explained (CVE); right panel: factor loadings. PC: principal component; bw: backward maximum; fw: forward maximum; abs: absolute score.

S4 Table. Priors for group-level parameters in the computational models.

| | | $\mu$ | $\sigma$ |
|---|---|---|---|
| DR models | $\alpha$ | $N(0, 10)$ | $U(0, 20)$ |
| | $\beta$ | $N(0, 10)$ ⊤ $[0, \infty)$ | $U(0, 20)$ |
| | $\varphi$ | $N(0, 10)$ | $U(0, 20)$ |
| | $\rho$ | $N(0, 10)$ | $U(0, 20)$ |
| KF models | $\beta$ | $N(0, 10)$ ⊤ $[0, \infty)$ | $U(0, 20)$ |
| | $\varphi$ | $N(0, 10)$ | $U(0, 20)$ |
| | $\rho$ | $N(0, 10)$ | $U(0, 20)$ |
| | $\alpha_{HOP}$ | $N(0, 10)$ | $U(0, 20)$ |
| | $\gamma$ | $N(0, 10)$ | $U(0, 20)$ |
| | $\beta_{drug\,effect}$ | $N(0, 10)$ | $N(0, 10)$ |
| | $\varphi_{drug\,effect}$ | $N(0, 10)$ | $N(0, 10)$ |
| | $\rho_{drug\,effect}$ | $N(0, 10)$ | $N(0, 10)$ |
| | $\alpha_{HOP\,drug\,effect}$ | $N(0, 10)$ | $N(0, 10)$ |
| | $\gamma_{drug\,effect}$ | $N(0, 10)$ | $N(0, 10)$ |

Note. Priors are specified in distributional form (e.g., N = normal, U = uniform), with truncation (⊤) where applicable. DR: Delta rule; KF: Kalman filter; β: random exploration (softmax inverse temperature); φ: directed exploration; ρ: perseveration strength; $\alpha_{HOP}$: habit update; γ: total uncertainty weight.

S6 Table. Descriptive statistics and Bayes' factors for drug condition differences (sample 1, *N* = 75) in model-agnostic measures.

| | Placebo | | L-DOPA | | | |
|---|---|---|---|---|---|---|
| | *M* | *SD* | *M* | *SD* | *BF* | |
| Response time | 0.45 | 0.08 | 0.47 | 0.09 | 22.09 (0+) | 0.05 (+0) |
| Total points | 12086 | 1036 | 12201 | 14972 | 7.69 (01) | 0.13 (10) |
| Stay | 0.60 | 0.17 | 0.61 | 0.16 | 5.30 (0-) | 0.19 (-0) |
| Switch | 0.40 | 0.17 | 0.39 | 0.16 | 5.21 (0+) | 0.19 (+0) |
| UNC-based exploration | 0.15 | 0.10 | 0.14 | 0.07 | 6.28 (0+) | 0.16 (+0) |
| Random exploration | 0.25 | 0.11 | 0.25 | 0.11 | 7.47 (01) | 0.13 (10) |
| Optimal choices | 0.60 | 0.09 | 0.60 | 0.09 | 7.74 (01) | 0.13 (10) |
| | Stay | | Switch | | | |
| | *M* | *SD* | *M* | *SD* | *BF* | |
| Response time | 0.46 | 0.09 | 0.45 | 0.08 | 1.32 (01) | 0.76 (10) |
| | UNC-based | | RAND | | | |
| | *M* | *SD* | *M* | *SD* | *BF* | |
| Response time | 0.45 | 0.08 | 0.46 | 0.08 | 22.15 (10) | 0.05 (01) |

Note. Stay and switch indicate repeating and changing the previous choice, respectively, uncertainty-based exploration reflects choosing the option not selected for the longest time, random exploration includes switch trials not classified as uncertainty-based, and optimal choices represent choosing the highest-value option. All measures are proportions per drug condition. Bayes' factor (BF) notation indicates evidence for the null versus alternative hypothesis: 0+/0– indicate evidence for the null in directional tests with positive/negative expected effects, +0/–0 indicate evidence for the alternative in directional tests with positive/negative expected effects, and 01/10 indicate evidence for the null or alternative, respectively, in two-sided tests. UNC-based: uncertainty-based exploration; RAND: random exploration

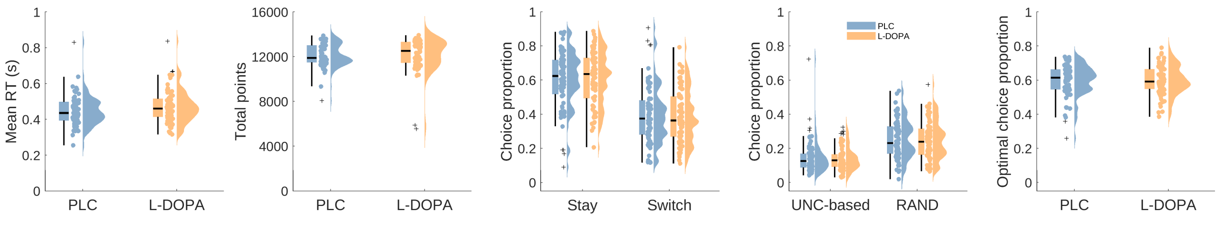


S7 Figure. Reaction times, total point scores and proportion of model-agnostic choice types, i.e. stay and switch decisions (repeating and changing the previous choice, respectively), uncertainty-based exploration (choosing the option not selected for the longest time), random exploration (switch trials not classified as uncertainty-based) and optimal choices (choosing the highest-value option), for the placebo (blue) and L-DOPA (orange) condition (sample 1, *N* = 75). The horizontal line depicts the median, crosses indicate outliers (values below Q1-1.5*IQR and above Q3+1.5*IQR). PLC: placebo; UNC-based: uncertainty-based exploration; RAND: random exploration.

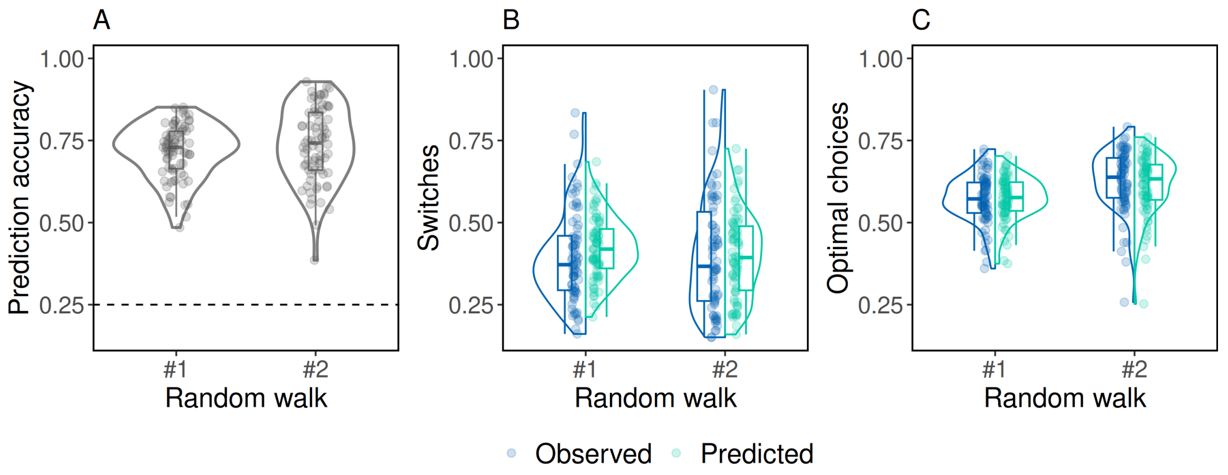


S8 Figure. Posterior predictive checks (sample 1, *N* = 75). Comparison of simulated to observed choices, separately for each random walk and averaged across simulations. Panel A: predictive accuracy as the proportion of correct choice predictions by the winning model (mode of option across samples for each trial and participant) compared to observed choices. Panel B: proportion of switches (choosing another option). Panel C: proportion of optimal choices (choosing the highest-value option). Horizontal dashed line: chance level. Box: Q1 to Q3 and median line. Whiskers: 1.5 × inter-quartile-range. Violin: density with Gaussian kernel. Points: single participants.

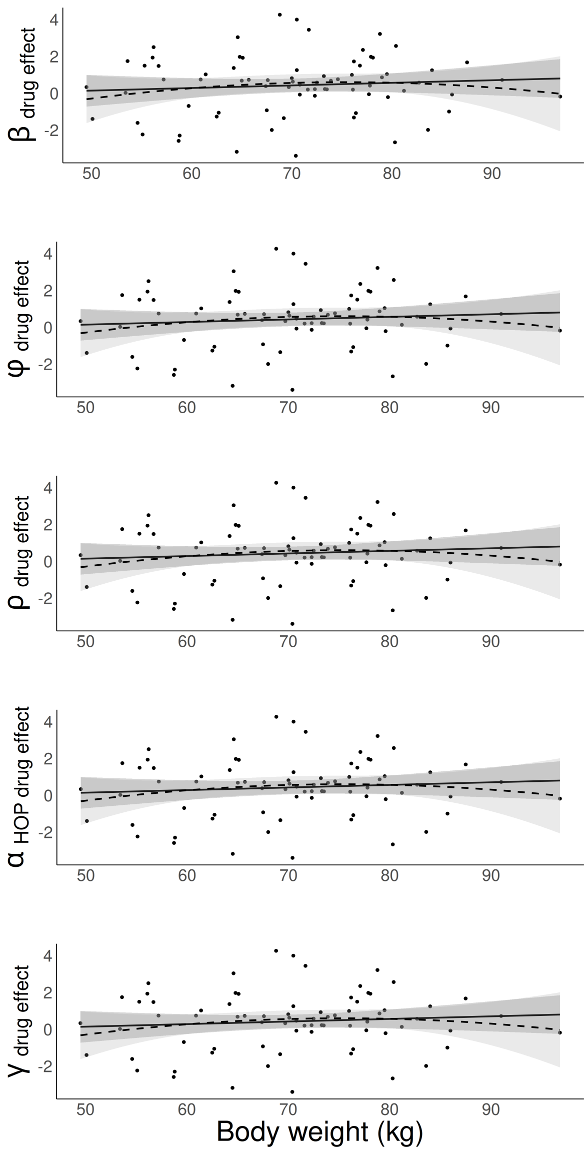


S9 Figure. Relationship between body weight (sample 1, $N$ = 75) and the drug effects (means of single-subject posterior distributions for the best-fitting Bayesian learner model including terms for directed (i.e., uncertainty-based) exploration, higher-order perseveration and total uncertainty ($SM_{+E\ +HOP\ +TU}$). Points: single-subject values; solid black line: regression line; grey ribbon: standard error. $\beta$: random exploration (softmax inverse temperature); $\varphi$: directed exploration; $\rho$: perseveration strength; $\alpha_{HOP}$: habit update; $\gamma$: total uncertainty weight.

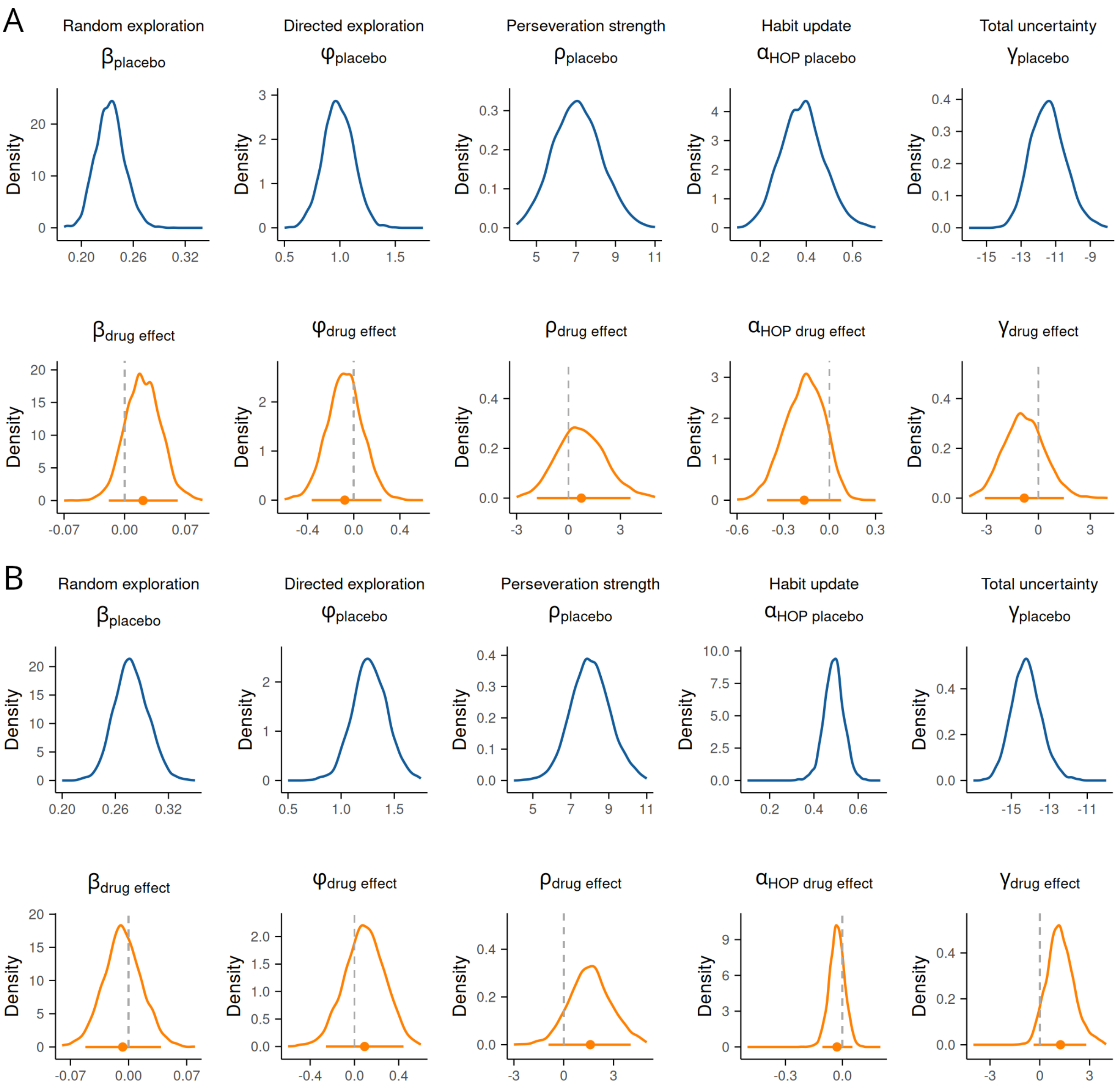


S10 Figure. Posterior distributions of the group-level parameter means for the best-fitting Bayesian learner model, which includes terms for random and directed exploration, higher-order perseveration, and total uncertainty ($SM_{+E\ +HOP\ +TU}$). The sample (sample 1, *N* = 75) was split by median body weight (*MD* = 70.8) into two subgroups: panel A: lower-weight participants (*n* = 39), panel B: higher-weight participants (*n* = 36). In each panel, the top row (blue distributions) represents parameters in the placebo condition modeled as baseline, and the bottom row (orange distributions) represents drug effects modeled as additive changes from placebo to L-DOPA. Vertical dashed line: $x=0$. Horizontal solid orange line: 95% highest posterior density interval. $\beta$: random exploration (softmax inverse temperature); $\varphi$: directed exploration; $\rho$: perseveration strength; $\alpha_{HOP}$: habit update; $\gamma$: total uncertainty weight.

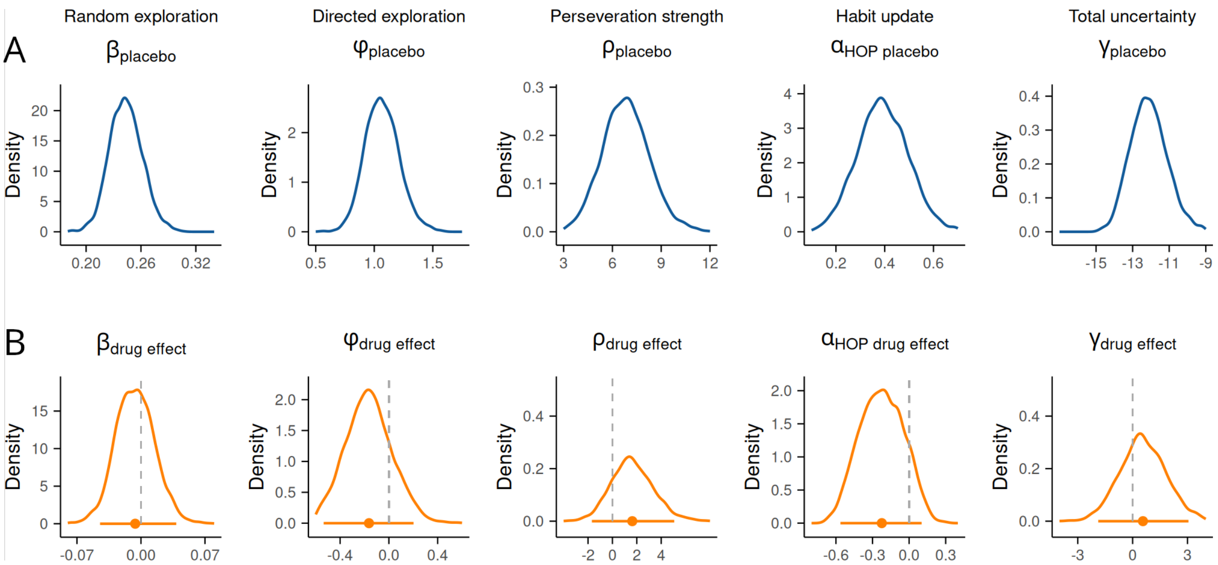


S11 Figure. Posterior distributions of the group-level parameter means of the female subsample of sample 1 (*n* = 32) for the best-fitting Bayesian learner model including terms for random and directed exploration, higher-order perseveration and total uncertainty ($SM_{+E +HOP +TU}$). Panel A (top row, blue distributions): parameters in placebo condition modelled as baseline. Panel B (bottom row, orange distributions): drug effects modelled as additive changes from placebo to L-DOPA. Vertical dashed line: $x = 0$. Horizontal solid orange line: 95% highest posterior density interval. $\beta$: random exploration (softmax inverse temperature); $\varphi$: directed exploration; $\rho$: perseveration strength; $\alpha_{HOP}$: habit update; $\gamma$: total uncertainty weight.

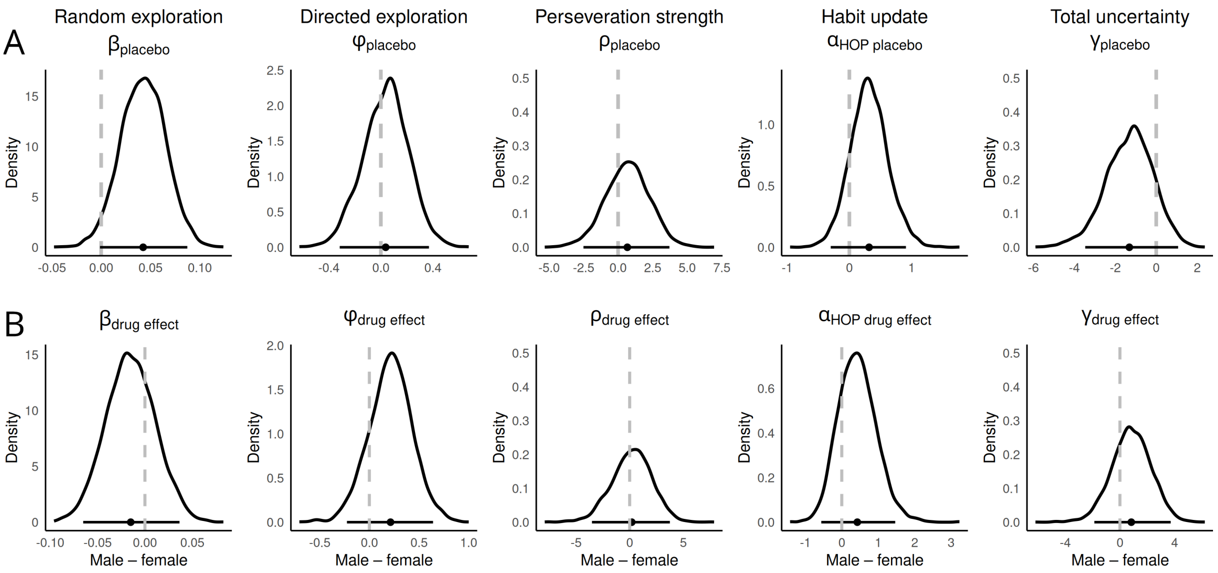


Figure S12. Differences between the posterior distributions of the group-level parameter means from the male (sample 2, *N* = 74) and female (sample 1, *n* = 32) groups, from the best-fitting Bayesian learner model, including terms for random and directed exploration, higher-order perseveration, and total uncertainty($SM_{+E\ +HOP\ +TU}$). Panel A (top row): parameters in placebo condition modelled as baseline. Panel B (bottom row): drug effects modelled as additive changes from placebo to L-DOPA. Vertical dashed line: $x=0$. Horizontal solid orange line: 95% highest posterior density interval. $\beta$: random exploration (softmax inverse temperature); $\varphi$: directed exploration; $\rho$: perseveration strength; $\alpha_{HOP}$: habit update; $\gamma$: total uncertainty weight.

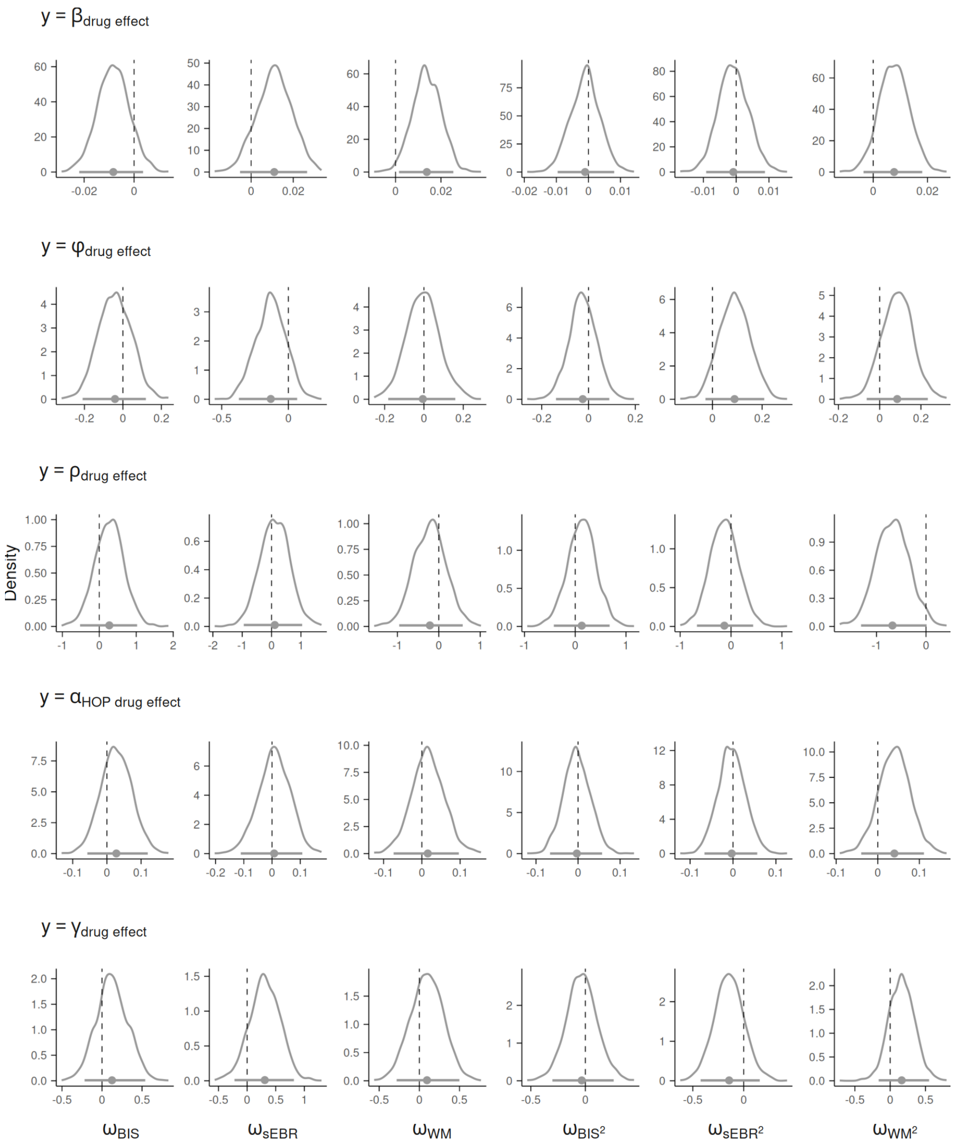


Figure 13. Posterior distributions of the linear (column 1 to 3) and quadratic (column 4 to 6) regression coefficients ($\omega$), regressing the drug effect parameters of the Bayesian learner model with $SM_{+E\ +HOP\ +TU}$ onto the (squared) putative DA proxy measures impulsivity (BIS-15 score), sEBR and WM capacity. Analyses shown here excluded two participants with WM capacity scores < 2 SDs below the sample mean. The posterior distribution of the intercept is omitted. Vertical dashed line: $x=0$. Horizontal solid line: 95% highest posterior density interval. $s$: drug effects modelled as additive changes from placebo to L-DOPA. $\beta$: random exploration (softmax inverse temperature); $\varphi$: directed exploration; $\rho$: perseveration strength; $\alpha_{HOP}$: habit update; $\gamma$: total uncertainty weight; BIS-15: scores for the Barratt Impulsiveness Scale (short form); sEBR: spontaneous eye blink rate, WM: working memory capacity.

S14 Table. Summary of generalised linear mixed-effects model results for effects of drug, uncertainty of the chosen option, and total (summed) uncertainty on the number of pre-choice fixation shifts (sample 1, *N* = 75).

| Phase | Effect | Estimate | *SE* | *t* | DF | *p* |
|---|---|---|---|---|---|---|
| Pre-choice | Intercept | -2.361 | 0.203 | -11.60 | 28356 | < .001 |
| | Drug | 0.237 | 0.256 | -0.93 | 28356 | 0.353 |
| | $\sigma_{chosen}$ | 0.071 | 0.013 | 5.38 | 28356 | < .001 |
| | Σσ | -0.032 | 0.006 | -5.44 | 28356 | < .001 |
| | Drug : $\sigma_{chosen}$ | -0.003 | 0.019 | -0.17 | 28356 | 0.868 |
| | Drug : Σσ | 0.006 | 0.008 | 0.78 | 28356 | 0.435 |

Note. Drug effect was coded with placebo as the baseline condition. The model was fitted using a Poisson distribution with a log link function. SE: standard error; DF: degrees of freedom; $\sigma_{chosen}$: uncertainty of chosen option; Σσ: total uncertainty.

S15 Table. Summary of generalised linear mixed-effects model results for effects of drug, trial, cognitive variables and choice type on the area under the curve (AUC) of the pupil dilation trajectory across different task phases (sample 1, *N* = 75).

| | GLME | | | | | | Posthoc | | |
|---|---|---|---|---|---|---|---|---|---|
| Phase | BIC | Effect | Estimate | SE | t | p | F | p | DF |
| Pre-choice | -26456 | Intercept | 0.878 | 0.005 | 185.66 | < .001 | | | 17204 |
| | | Drug | 0.010 | 0.003 | 2.98 | .003 | | | |
| | | Trial | 0.000 | 0.000 | -3.87 | < .001 | | | |
| | | Drug : trial | 0.000 | 0.000 | 0.20 | .839 | | | |
| | | EV | -0.007 | 0.001 | -6.02 | < .001 | | | |
| | | $\sigma_{chosen}$ | -0.008 | 0.001 | -6.46 | < .001 | | | |
| | | Σσ | -0.005 | 0.001 | -4.28 | < .001 | | | |
| | | Choice type$_{directed}$ | 0.008 | 0.004 | 1.92 | .054 | | | |
| | | Choice type$_{random}$ | 0.012 | 0.003 | 3.96 | < .001 | | | |
| | | Directed vs. random | | | | | 1.10 | .294 | 1,17204 |
| Feedback | -24846 | Intercept | 0.955 | 0.004 | 273.73 | < .001 | | | 17202 |
| | | Drug | 0.010 | 0.003 | 3.04 | .002 | | | |
| | | Trial | 0.000 | 0.000 | -1.53 | .126 | | | |
| | | Drug : trial | 0.000 | 0.000 | -1.58 | .114 | | | |
| | | EV | -0.007 | 0.001 | -5.93 | < .001 | | | |
| | | $\sigma_{chosen}$ | -0.006 | 0.001 | -4.13 | < .001 | | | |
| | | Σσ | -0.004 | 0.001 | -3.81 | < .001 | | | |
| | | PE$_{signed}$ | 0.002 | 0.001 | 2.43 | .015 | | | |
| | | PE$_{unsigned}$ | 0.007 | 0.001 | 7.49 | < .001 | | | |
| | | Choice type$_{directed}$ | -0.002 | 0.004 | -0.48 | .634 | | | |
| | | Choice type$_{random}$ | 0.006 | 0.003 | 1.82 | .069 | | | |
| | | Directed vs. random | | | | | 4.21 | .040 | 1,17202 |
| ITI | -22005 | Intercept | 1.455 | 0.011 | 138.82 | < .001 | | | 17134 |
| | | Drug | 0.006 | 0.004 | 1.61 | .108 | | | |
| | | Trial | 0.000 | 0.000 | -5.28 | < .001 | | | |
| | | Drug : trial | 0.000 | 0.000 | 0.82 | .413 | | | |
| | | EV | -0.005 | 0.001 | -3.32 | .001 | | | |
| | | $\sigma_{chosen}$ | -0.005 | 0.002 | -3.24 | .001 | | | |
| | | Σσ | -0.007 | 0.001 | -5.81 | < .001 | | | |
| | | PE$_{signed}$ | 0.002 | 0.001 | 2.06 | .039 | | | |
| | | PE$_{unsigned}$ | 0.010 | 0.001 | 9.40 | < .001 | | | |
| | | Choice type$_{directed}$ | 0.004 | 0.005 | 0.90 | .368 | | | |
| | | Choice type$_{random}$ | | | | .001 | | | |
| | | Directed vs. random | | | | | 3.27 | .071 | 1,17134 |

Note. Drug effect was coded with placebo as the baseline condition; choice type was coded with exploitation as the baseline. Models were fitted using an identity link function. Post-hoc pairwise contrasts were conducted for choice type directed vs. random exploration using MATLAB's coefTest function. All continuous predictors (i.e., uncertainty measures and decision variables) were standardised. BIC: Bayesian information criterion; SE: standard error; DF: degrees of freedom; EV: expected value; $\sigma_{chosen}$: uncertainty of chosen option; Σσ: total uncertainty; PE: prediction error; ITI: inter-trial-interval.
